# Built-in integrated living electronics: from biosynthesis to modulation of neuronal function

**DOI:** 10.64898/2026.03.12.710354

**Authors:** Giuseppina Tommasini, Marika Iencharelli, Silvia Santillo, Philip Schaefer, Daniela Intartaglia, Martina Blasio, Graziano Preziosi, Maria Antonietta Ferrara, Gennaro Sanità, Emanuela Esposito, Giuseppe Coppola, Mattia Zangoli, Francesca Di Maria, Angela Tino, María Moros, Claudia Tortiglione

## Abstract

Neuroelectronic interfaces hold great promise to restore functions in neurological disorders or motor dysfunctions, but current devices struggle to integrate seamlessly within living tissues. Here we report a transformative approach to create bionic neurons that autonomously build integrated fluorescent fibrils and demonstrate their role as neuromodulators. Using a combination of cell biology, ultrastructural, imaging and nanospectroscopical approaches, we deciphered the unique biosynthetic pathway employed by the cells to self-fabricate these nanoelectronics and uncover their hybrid structure. Importantly, patch clamp recordings revealed their neuromodulatory potential, through the perturbation of membrane electrical properties and the early rising phase of the action potential. Deciphering how basic molecular elements self-organize into complex architectures within biological environments could unlock the ability to engineer natural electroactive systems directly inside living organisms. This capability could be used to create conductive pathways between arbitrarily defined neurons, microcircuits, or nervous system regions, effectively writing connections into living brains.

## Introduction

Neuroelectronic interfaces enable the transfer of signals between the nervous system and an external device, facilitating the investigation of fundamental neuroscience questions, but also regenerative issues, bioengineering needs and clinical issues. Conventional electrodes transduce cellular activity into electronic information (recording) and transmit signals into tissue modulating (by stimulation) the communication between cells. Through varying levels of invasiveness, several biomedical devices have been developed to access different forms of neural information, from implantable devices to engineered tissues ^[1]^: ultrathin flexible electronics replaced the first generation of stiff and non-compliant neural probes ^[2]^; stretchable mesh nanoelectronics allowed to dynamically track electrical activity during tissue differentiation (human stem cell-derived cardiac tissue and pancreatic islets) and brain development ^[3]^; bioinspired neuron-like electrodes allowed interrogation of individual neurons into brain parenchyma ^[4]^; bioresorbable and transient electronics, obviated potential adverse effects of chronic implants ^[5]^. Despite giant advances, and the deployment of many devices in the clinical environment, there are still challenges to bridge the gap between the full potential of neuroelectronic interfaces and their translation into clinical practice. Major challenges include the methods for perfectly and naturally integrating the electronic component into the tissue, the detrimental reduction/oxidation reactions at the biotic/abiotic interface, the foreign body response, especially for long-term use, and the lack of adaptability to the unique characteristics of the biological environment they interact with, in terms of biochemistry, topography, and electrical properties, which demands electrodes customized for each tissue.

To address some of these issues, the *in situ* cell-mediated fabrication of conductive devices represent a revolutionary approach for the seamless integration of electronic components into living systems ^[6]^, coevolving together and becoming functionally inseparable. Recently, organic mixed ionic-electronic conductors (OMIECs), capable of simultaneously transport both electronic and ionic species, have been proposed as promising candidates for *in situ* chemical assembly of conductive interfaces ^[7]^. Pioneering works from Stavrinidou et al. showed that the water soluble thiophene-based monomer ETE-S (2,5-bis(2,3-dihydrothieno[3,4-b][1,4]dioxin-5-yl)thiophene) undergoes polymerization in plants due to the endogenous peroxidase activity ^[8]^. Later, ETE-S polymerization was achieved in a living invertebrate, *Hydra vulgaris* ^[9]^. Beside revealing the neuromodulatory effect of the precursor monomer ^[10]^, the small polyp was shown able to fabricate electrically conductive interfaces in specific cells expressing peroxidase activity, fully integrated and evolving with the polyp life cycle. To improve spatial control of polymerization, alternative approaches based on chemical or genetic engineering have been developed. For instance, the injection of a gel containing peroxidase enzymes, peroxides and organic monomers into leech neural tissue promoted the metabolite-induced formation of conductive polymer gels, that lowered electrode-nerve impedance and improved stimulation performance ^[11]^. Layered patterns of conductive polymers were generated *ex vivo* in zebrafish fins using thiophene-based monomers and photocatalysts, although photocatalyst toxicity remains a limitation ^[12]^. To address this, similar EDOT-based monomers were employed to drive light controlled polymerization of conductive structures with defined morphology on mouse skin^[13]^. As alternative, genetically targeted photosensitizers were employed to polymerize conductive polyaniline in neurons with precise spatiotemporal control, enabling stepwise tuning of membrane capacitance and intrinsic excitability in neurons and freely moving *Caenorhabditis elegans*^[14]^. Together, these advances in chemical, genetic and optogenetic polymerization strategies highlight the need for interfacial building blocks that can be efficiently delivered into the host cells or pass through biological barriers and self-organize into conductive structures, endowing the host with novel “cyborg” properties. To this end, the small semiconductive oligothiophene, 3,5-dimethyl-2,3’-bis(phenyl)dithieno[3,2-b;2’,3’-d]thiophene-4,4-dioxide (DTTO), stands out thanks to its capability to assemble into fluorescent and conducting microfibrils ^[15]^. When living cells or *Hydra* tissues are exposed to micromolar doses of DTTO, highly fluorescent microfibrils of spectacular shapes form spontaneously, lying embedded into tissues, interconnecting adjacent cells, or meandering outside the tissue ^[16]^. These microfibrils feature properties profoundly different from DTTO monomer aggregates formed in certain solvents or in solid-state conditions ^[17]^. The fact that DTTO fibril formation relies on intracellular mechanisms of living cells that cannot be replicated outside them raises significant questions, both on the mechanisms underlying the supramolecular assembly of DTTO within the cell environment and on their effect on the cell physiology.

Here we investigated how DTTO dose influences fibril formation, identified its internalization route and intracellular trafficking and demonstrated, through multimodal analysis, its sequestration into lipid droplets within the lipid metabolism pathway. Remarkably, autonomous DTTO self-assembly within these droplets yields biohybrid fibrils featuring a crystalline DTTO core encased in a proteinaceous shell, as revealed by nanospectroscopy. Intertwined with the cytoskeleton, these fibrils exert a functional role in host cells, as shown by electrophysiological recordings: they act as living conductive interfaces that increase membrane capacitance, depolarize the resting potential, and modulate the early rising phase of the action potential, thereby serving as intrinsic neuromodulatory electrodes. These results introduce a strategy for the cell-primed construction of integrated nanoelectronics that modulate neuronal excitability directly from within.

## RESULTS

### Autonomous production and seamless integration of fluorescent fibrils in neuronal cells

A typical fibril fully embedded into human-derived neuroblastoma cells (SH-SY5Y), used as model of neurons, is shown in Fig. 1A. Cells were stained with anti-βtubulin antibody to better map the fibril spatial distribution. Disparate and unique fibril geometries were crafted, fully integrated within the cell architecture, sometime even crossing the nucleus (Fig. S1, Movie S1-S2).

**Fig. 1.**
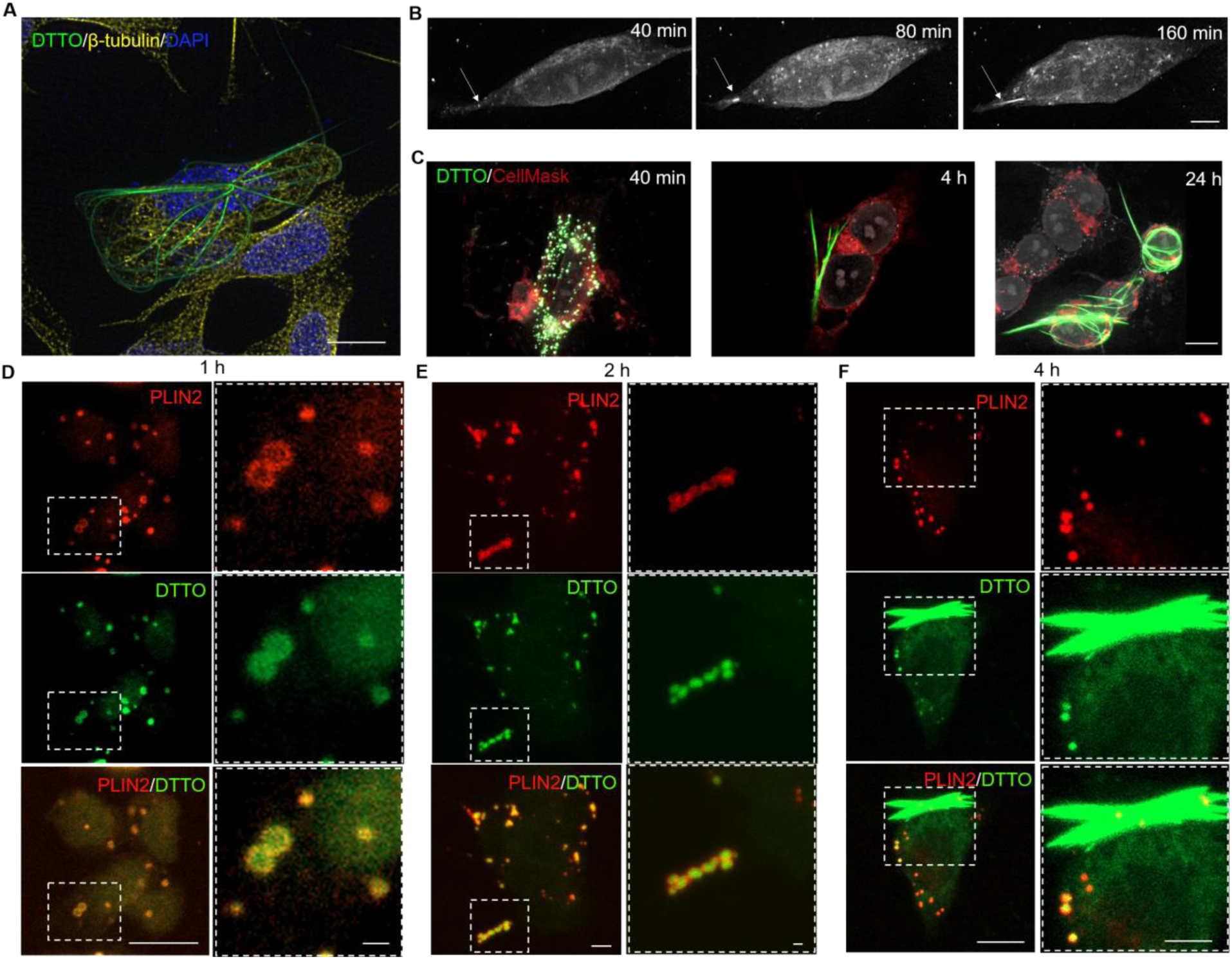
Real-time dynamic of fibril production and DTTO confinement in lipid droplets. **(A)** Super-resolution imaging of SH-SY5Y cells treated 24 hours with DTTO 5 µg/ml, showing a DTTO fibril (green), β-tubulin (yellow), and nuclei stained with DAPI (blue). Scale bar: 10 µm. **(B)** Time-course holotomography images acquired at 40 min, 80 min and 160 min. Both monomer (white spots) and fibril (white line) are visible due to DTTO refractive index (RI= 1.36-1.38). Scale bar: 5 µm. **(C)** Merged fluorescence and holotomography images of SH-SY5Y cells treated with DTTO (green) for 40 min, 4 hours and 24 hours and stained with CellMask (red, labelling plasma membrane lipids), before fixation. A single Z-plane of representative cells for each time is shown. Nuclei are shown in grey (RI=1.362-1.367). Fibrils fully embedded onto the cells are clearly visible through all the Z planes (Movies S6-S8). Scale bar: 10 µm. **(D-F)** Representative images of PLIN2 (red)-DTTO (green) colocalization in SH-SY5Y cells treated with DTTO 5 µg/ml for 1, 2 and 4 hours. The dashed white boxes show magnified insets of the selected regions. Scale bars: main 10 µm, inset 1 µm.

A systematic quantitative analysis was performed to evaluate DTTO effect on cell viability and fibril yield. A transient cytotoxicity was observed at the highest dose (Fig. S2A-B, D), while low dose (5 µg/mL) did not affect cell viability, even after long-term exposure (Fig. S2C, E). Fibril yield increased with both DTTO concentration and incubation time: high dose rapidly induced fibril formation in a minority of cells, whereas a prolonged low-dose exposure ultimately produced an even higher fraction of fibril-containing cells (Fig. S3). DTTO uptake, estimated by fluorescence microscopy and flow cytometry, appeared rapid, dose- and time-dependent, with almost all cells internalizing the monomer within the first hour and the uptake rising steeply during the initial 20 min (Fig. S4, S5). Remarkably, fibril morphology was found independent of DTTO dose (Fig. S6A), while fibril length and area grew with incubation time, consistent with an active, time-dependent elongation process (Fig. S6B).

The dynamic of the fibril growth was studied in real-time by holotomography (HT), a label-free, non-invasive, quantitative imaging technique based on the measurement of refractive index (RI) distributions within the cell ^[18]^. Fig. 1B and Fig. S7 show the 3D RI tomogram of living neuroblastoma cells, reconstructed from multiple 2D acquired holograms, up to 160 min of continuous incubation with DTTO, clearly showing the initiation of the fibril formation. Next, the fibril intricate interaction with the cell structure was investigated by using a membrane marker (Fig. 1C Movie S3-S5). At 40 min, most of the DTTO signal appears as punctuated fluorescence throughout the cell, while at 4 and 24 hours, fibrils, initially confined within the cell, were observed progressively growing, twisting and creating bundles also outside the cell membrane, without compromising the cell viability.

### DTTO internalization route and intracellular trafficking

In order to evaluate the DTTO uptake mechanism, SH-SY5Y cells underwent a series of treatments targeting passive and active internalization pathways. DTTO treatment performed at 4 °C caused a marked reduction in intracellular DTTO fluorescence, as shown by fluorescence microscopy (Fig. S8A-B) and flow cytometry (Fig. S8C-E), with no effect on cell morphology and viability (Fig. S8E). These data suggest that while DTTO can still associate to the cell membrane at low temperature, its energy-dependent internalization is largely impaired, consistent with a requirement for membrane fluidity and/or active endocytic processes. Immunofluorescence assays performed to test clathrin-mediated endocytosis revealed no colocalization between DTTO-containing vesicles and clathrin signals, supporting a clathrin-independent uptake of DTTO rather than through a classical endocytic pathway within this temporal window (Fig. S9). This was further confirmed by the finding that the impairment of endosomal trafficking by ammonium chloride (NH₄Cl), does not affect DTTO internalization ^[19]^(Fig. S10). Altogether these results demonstrate that DTTO internalization in SH-SY5Y cells relies on both passive diffusion and active processes. The significant decrease in uptake at 4 °C may depend on reduced membrane fluidity and inhibition of energy-dependent processes, while the residual fluorescence signal arises from passive diffusion.

### DTTO is sequestered in lipid droplets and enter the lipid pathway

A key question was whether internalized DTTO is targeted to specific organelles. Data obtained from HT strongly support the hypothesis that lipid droplets (LDs) act as the principal organelles involved in the intracellular sequestration of DTTO, as indicated by their closely matching refractive indices (mean RI=1,38)(Fig. 1B-C) ^[20]^. Correlative 3D fluorescence imaging with the LD marker BODIPY (boron-dipyrromethene) confirms this localization, showing DTTO signal closely overlapping with BODIPY as early as 5 min after incubation (Fig. S11). Quantitative analysis over the first 30 min of incubation demonstrated that the rapid sequestration of DTTO into LDs takes place continuously as more DTTO reaches the cytoplasm (Fig. S11). LDs are dynamic organelles primarily recognized for their role in lipid storage and energy homeostasis. They consist of a hydrophobic core of triacylglycerols and sterol esters, surrounded by a phospholipid monolayer decorated with cholesterol and specific proteins, such as perilipins. These proteins regulate lipolytic activity on the LD surface and participate in LD biogenesis ^[21]^. Immunofluorescence assays revealed robust colocalization of perilipin-2 (PLIN2) and DTTO monomer before, during, and after fibril formation (Fig. 1D–F, Fig. S12). Changing the experimental conditions, *i.e.* higher DTTO dose and short incubation time, produced same results (Fig. S13), confirming that DTTO is sequestered within individual cytoplasmic LDs. The PLIN2 staining consistently observed within single LDs favors the hypothesis of a local confinement priming the DTTO nucleation and subsequent fibril assembly (Fig. 1F). However, the increase of the LD size detected over time (Fig. S13D-E) cannot rule out the possibility of a concomitant LD coalescence phenomenon mediating fibril growth. Of note, the almost complete absence of colocalization between DTTO and lysotracker (lysosomal marker) fluorescence signals excludes the lysosomal accumulation and/or degradation as trigger of the fibril growth (Fig. S14), while reinforcing the LD involvement.

### Fibril formation is mediated by an autophagy-based mechanism

To increase the LD content and favour fibril formation, lipid autophagy, one of the pathways for LD degradation in neuronal cells ^[22]^ was pharmacologically and physically modulated. Autophagy proceeds *via* phagophore formation, autophagosome-lysosome fusion, and cargo degradation by acid hydrolases ^[22a]^. The dependence of the autophagy process on cell’s physiological state could explain why only a subset of cells forms fibrils. To probe its role in mediating fibril formation the following approaches were used: *i*) cold shock at 4°C, to inhibit cellular metabolism and autophagic flux by reducing enzyme activity and ATP production ^[23]^; *ii*) serum deprivation, to enhance autophagy and promote autophagosome accumulation ^[24]^ and *iii*) NH₄Cl and bafilomycin A1 (BafA1) to block lysosomal acidification, autophagosome-lysosome fusion and degradation of autolysosome content ^[25]^. Changes in fibril yield were used as readout of how autophagy impacts on fibril biogenesis. When SH-SY5Y cells were treated with DTTO at 4°C, fibril formation was almost completely abolished, indicating the requirement of an active metabolism to favor not only the DTTO internalization but also the fibril synthesis (Fig. 2A, C). For instance, a brief cold pulse after DTTO treatment (37°C→4°C) also reduced fibril yield (from 11.5% to 4.3%), further supporting the need for active cellular processes during the early phase of fibril assembly. Serum deprivation, which enhances autophagy, increased the fibril yield by up to 30% compared to the standard DTTO treatment, whereas maintaining the serum before and during DTTO exposure significantly reduced fibril formation (by up to 20%, Fig. 2B, D). Consistently, blocking autophagic degradation with BafA1 or NH₄Cl further elevated fibril yield (Fig. 2E-F), confirming the crucial role of autophagy as key modulator during biofabrication of DTTO fibrils in SH-SY5Y cells.

**Fig 2.**
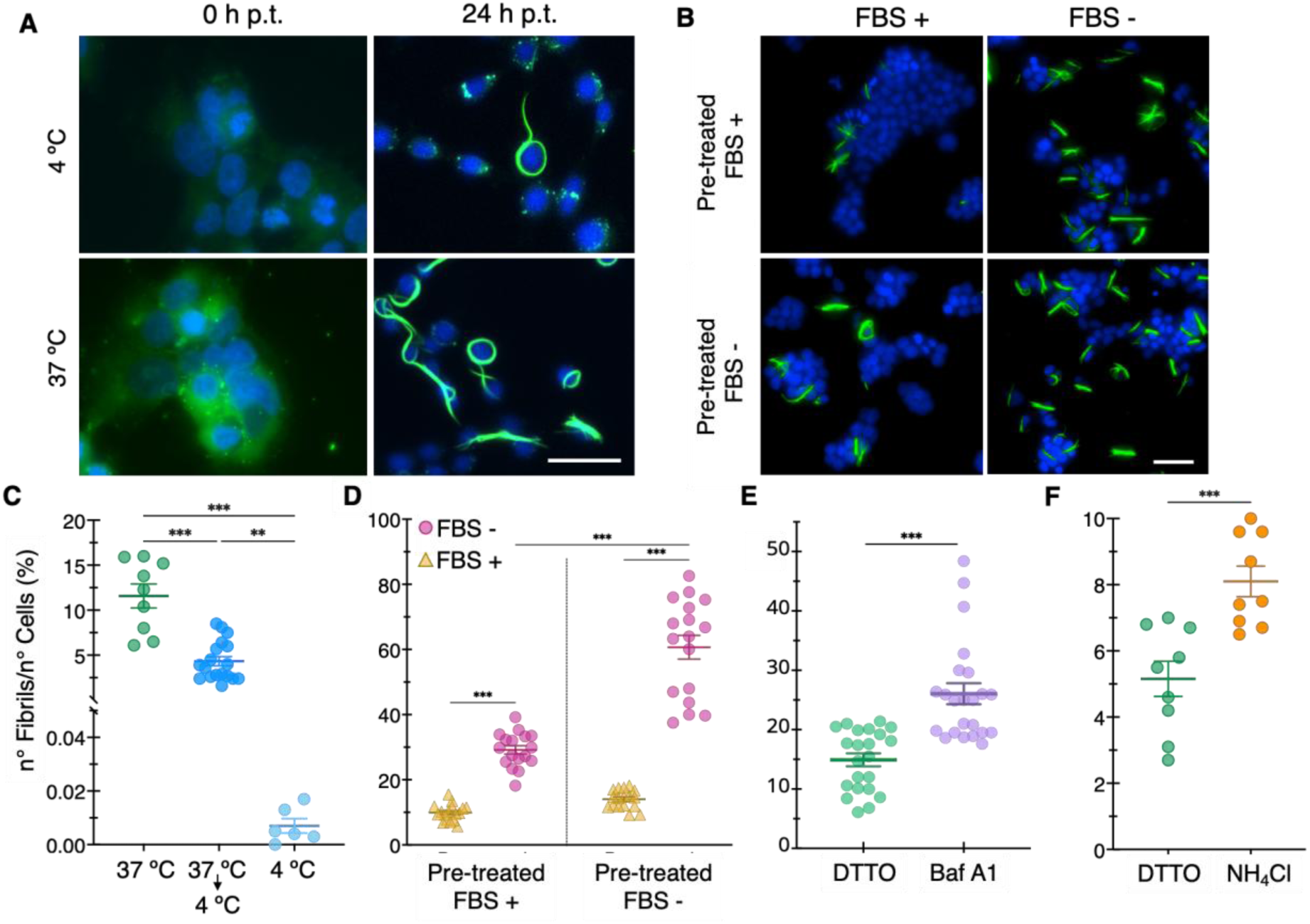
Active metabolism and autophagy are involved in the fibril biogenesis. **(A)** Representative images of SH-SY5Y cells treated with DTTO, either at 4 °C or 37 °C for 30 min. Images were acquired immediately post treatment (0 hours p.t.) and after 24 hours p.t. Scale bars: 25 µm. **(B)** Representative images of the four conditions analyzed in the serum deprivation assay. Cells were pre-treated for 24 hours in serum-containing or serum-free medium, and then treated with DTTO for 30 min under the same conditions. Scale bars: 50 µm. **(C-F)** Percentage yield of fibril production following cold shock **(C)**, serum availability **(D)**, bafilomycin-mediated **(E)**, and ammonium chloride-mediated autophagy inhibition **(F)**, calculated as explained in Methods section. Horizontal lines represent the mean ± SEM. Statistical analysis was performed using one-way ANOVA with Tukey’s post hoc test **(C)**, two-way ANOVA with Šídák’s post hoc test **(D)**, and Student *t*-test **(E–F)**; ** *p* < 0.01, *** *p* < 0.001.

### Ultrastructural analysis revealed biohybrid composition and complex morphological structure of DTTO fibrils

A crucial question is whether DTTO fibrils arise as pure crystalline assemblies or as biohybrids incorporating cellular proteins. Previous biochemical studies supported a co-assembly model ^[16a, 16b]^, identifying cell-specific proteins associated with DTTO fibrils, *i.e.*, collagen in fibroblasts and vimentin in neuroblastoma cells ^[15–16]^. A recent study proposed a “biologically assisted” polymorphic structure with unique DTTO molecular packing and protein-like material adsorbed post synthesis rather than intrinsically incorporated ^[17]^. In all cases the analyses were performed on fibrils extracted from the cells, raising concerns about potential artifacts due to certain detergents or extraction conditions. To solve these discrepancies, DTTO fibrils were analysed in their native cellular context. Immunofluorescence coupled to quantitative colocalization analysis using Pearson’s correlation coefficients (PCC) showed that fibrils moderately associate with β-tubulin (PCC= 0.626), weakly with α-tubulin (PCC= 0.421) and actin (PCC= 0.166) (Fig. S15-S16) while display strong correlation with vimentin (PCC= 0.775), indicating a strict interaction of DTTO fibrils with intermediate filament proteins. Z-stacks (Fig. S17-18) and confocal (Fig. S19) imaging further supported the involvement of the cytoskeleton filament network, acting in LD movement and clustering, as scaffold guiding the assembling of the biohybrid fibril structure.

To deeply investigate on the intrinsic incorporation of proteins within the fibril structure, a purification protocol using non-denaturing conditions was developed to preserve the fibril native structure (see Methods section). Conventional and cryogenic transmission electron microscopy (TEM and cryoTEM) revealed characteristic fibrillar morphologies, including both helicoidal and straight shapes (Fig. 3A-B). Planar fibrils displayed larger widths (1-2 µm) compared to thinner helicoidal fibrils, which averaged around 400 nm, in accordance with the size range observed in other cell lines by atomic force microscopy ^[15, 16c]^. A homogeneous electron dense area was detected in the inner part of the fibril, outlined by two regions of minor intensity which may correspond to an inner crystalline phase surrounded by other material. Notably, individual DTTO fibrils appear composed by multiple nanosized protofilaments (Fig. 3B, inset), suggesting a supramolecular assembly process that drives the formation of high-ordered fibrillar structures within the cellular environment. By coupling TEM with EDS (energy-dispersive X-ray spectroscopy) both sulphur, an integral element of the DTTO molecule, and nitrogen, which is characteristic of proteins and absent in the DTTO molecule, were detected within the fibril (Fig. 3C-E). Consistent with TEM, sulphur signal closely matched the fibril morphology, while nitrogen was detected also outside the fibril, albeit unevenly distributed (Fig. 3C-D). Longitudinal intensity profiling revealed that nitrogen was highly concentrated along the fibril edges (Fig 3D, F).

**Fig. 3.**
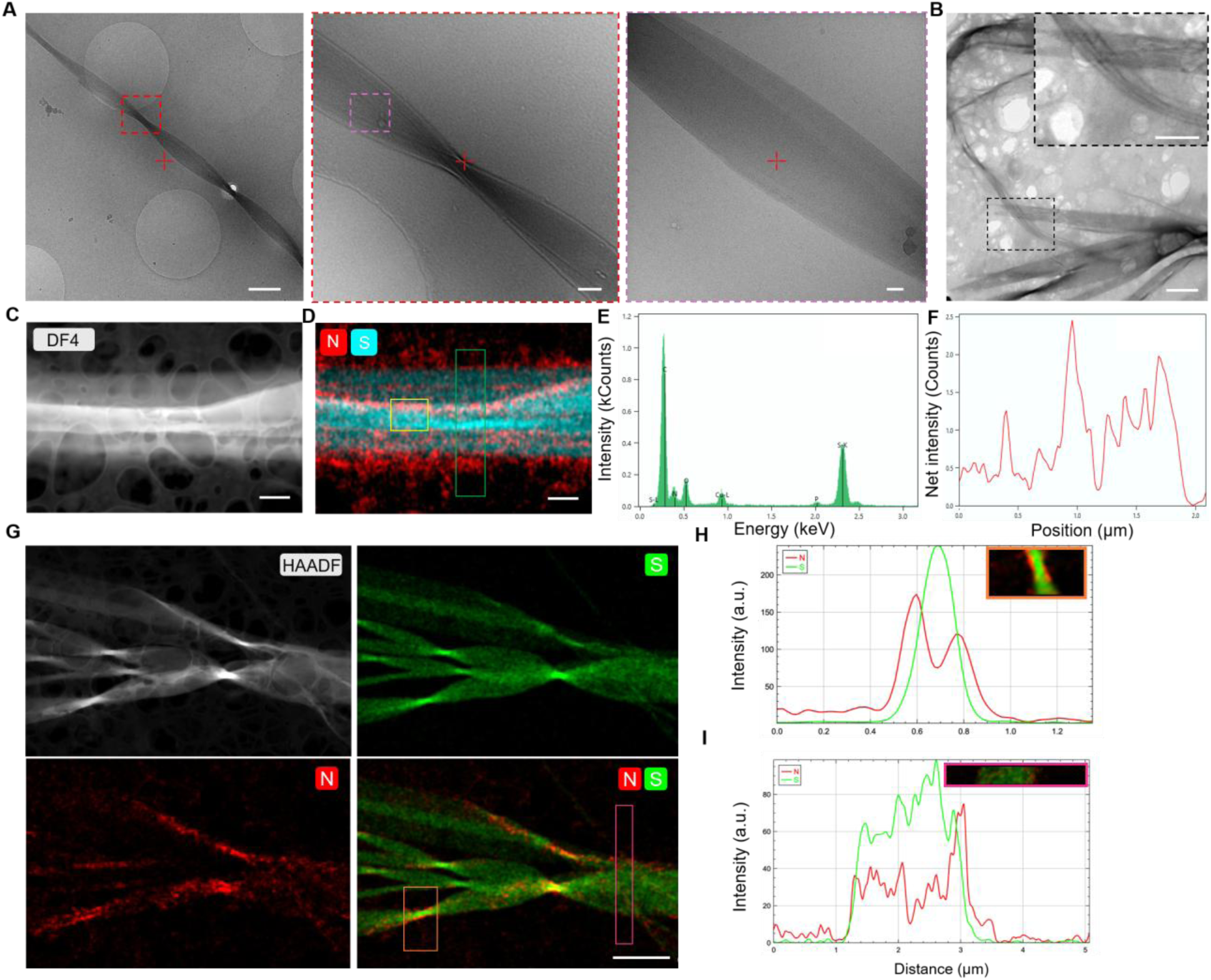
Ultrastructural and spectroscopical analyses reveal nitrogen on DTTO fibril surface. **(A)** CryoTEM images of cell-purified fibrils. Red and purple boxes show magnifications of the selected areas. Scale bars: 400 nm, 50 nm, 20 nm. **(B)** Fibrils isolated from SH-SY5Y cells showing characteristic helicoidal or straight shapes. Scale bars: 2 μm, inset 1 μm. **(C-F)** STEM-EDS characterization of DTTO fibrils. Fibrils were analysed by electron microscopy with Talos F200X G2 TEM/STEM operated at 200 kV. The images were first acquired with **(C)** a dark-field detector (DF4) and then analysed by EDS. Scale bars: 500 nm. **(D)** Elemental map of a single fibril showing the sulphur (cyan) and the nitrogen (red) signals. Scale bar: 500 nm. **(E)** Graph showing EDS spectrum obtained by a specific area (yellow box) and **(F)** the nitrogen distribution profile of the area included in the green box. **(G-I)** TEM/STEM EELS characterization of DTTO fibrils by TEM/STEM FEI Cubed Titan Themis operated at 300 kV. **(G)** Images were taken by High-Angle Annular Dark Field (HAADF) detector and then elemental maps of sulphur (green), and nitrogen (red), or both elements, were obtained by EELS. Scale bar: 2 μm. Fluorescence profile plots of the orange (**H**) and purple (**I**) areas were obtained by Fiji.

Nitrogen-enriched regions were frequently detected in empty spaces within the protofilaments, perhaps contributing to fibril filament cohesion (Fig. S20A). Additionally, nitrogen was concentrated at helical pitches of the fibrils, particularly along the edges, forming well-defined channels for fibril insertion, as confirmed by longitudinal intensities plots (Fig. S20 B-C).

Pushing further the resolution and sensitivity to light elements such as nitrogen, TEM coupled with electron energy loss spectroscopy (EELS) was employed. The elemental mapping revealed minimal nitrogen presence in regions adjacent to the fibril, with a consistent accumulation at the helical pitch and flanking areas (Fig 3G-I). To exclude that this mapping might reflect variations in fibril thickness, a nanoscale Fourier-transform infrared spectroscopy (nano-FTIR), coupling s-SNOM (scattering-type Scanning Near-field Optical Microscopy) with broadband illumination and FTIR-based detection, was employed to compare DTTO fibrils and DTTO aggregates formed spontaneously in DMSO. Nano-FTIR spectra from DTTO aggregates show characteristic DTTO molecular vibrational bands at 1143 and 1302 cm⁻¹, which were also present in all spectra from cell-derived fibrils (Fig. 4A-D, Fig. S21). Additionally, the latter exhibited amide I (1610–1700 cm⁻¹) and amide II (1490–1580 cm⁻¹) bands, indicative of proteins, throughout the fibril length (Fig. S21D-E). An additional peak around 1263 cm⁻¹ suggested a third component, potentially related to aromatic or amide C–N stretching or random coil protein structures (Fig. S21D-E). By mechanical amplitude imaging (Fig. 4B, Fig. S22A-B), a fibril with a rough region and a flatter area was serendipitously detected. Nano-FTIR mapping on both areas (Fig. S22) detected amide bands only on the rough area (Fig. 4D, Fig. S22) indicating a mechanical break-off of the protein coverage, possibly during sample preparation, and consequently the exposure of the DTTO-crystalline surface.

**Fig. 4.**
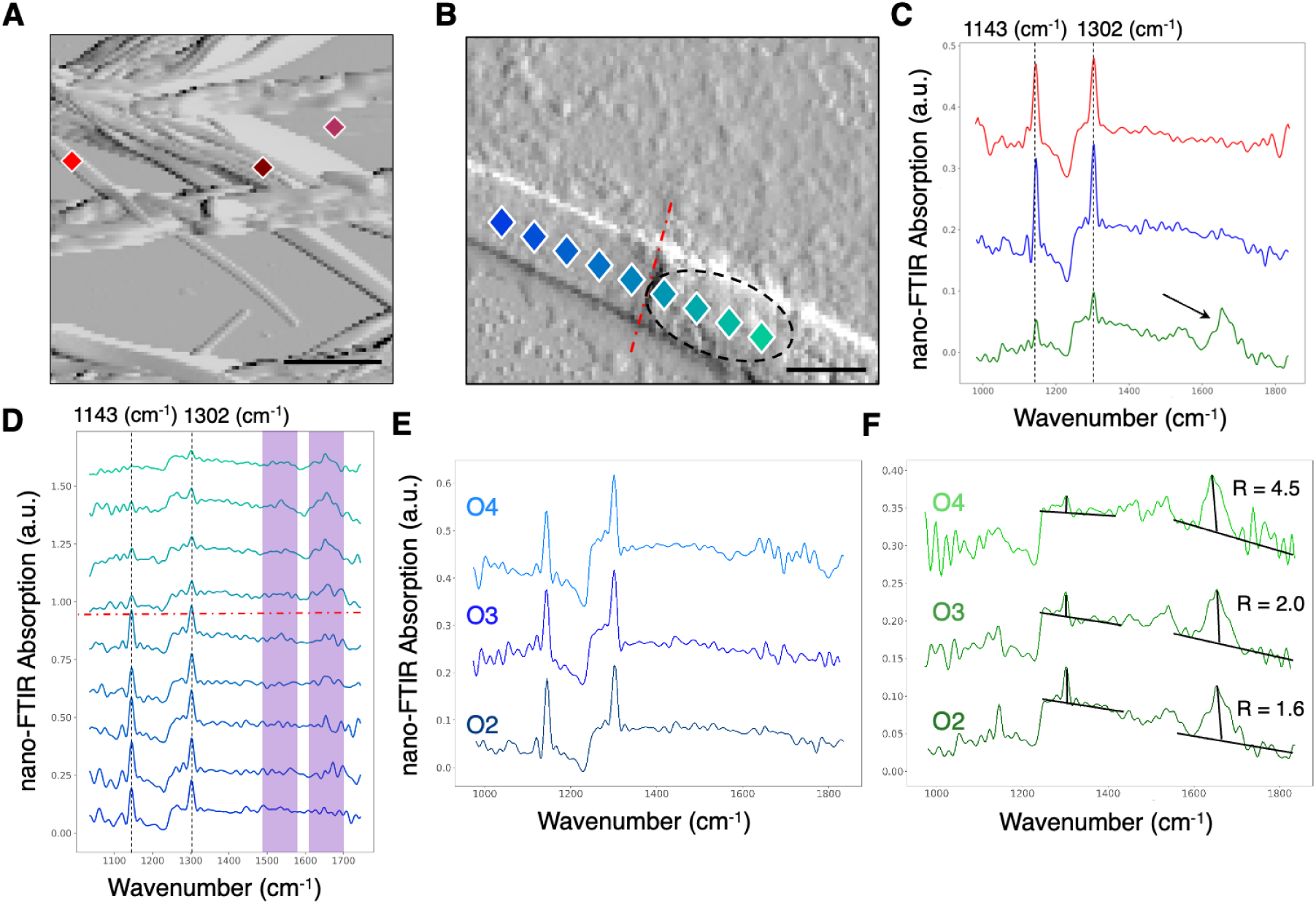
Nano-FTIR spectroscopy revealed heterogeneous DTTO fibril composition. **(A)** Phase images of synthetic fibrils, DMSO-aggregated and a cell-derived DTTO fibril **(B)** with heterogeneous composition, analysed by nano-FTIR spectroscopy. Each square indicates a specific area from which spectra were collected. Scale bars: 1 µm. In **(B)** the rupture point is shown by the dashed red line, while the black ellipse indicates the areas from which amide signals were detected. **(C)** Comparison between nano-FTIR spectra derived from synthetic fibril (**A**, red spectrum), cell-derived fibril without protein material (left from red dashed line in **B**, blue spectrum) and cell-derived fibril with protein material (black ellipse in **B**, green spectrum). More spectra on and next to the fibril are shown in Fig. S22. The characteristic sharp peaks of DTTO molecule at 1143 and 1302 cm^-1^ are present in all spectra, while the amide I and II bands are detected only in the green spectrum (arrow indicates the amide I). In **(D)**, spectra collected from each point shown in **B**. The light purple boxes highlight the amide bands. **(E-F)** Nano-FTIR spectra of 2^nd^ (O2), 3^rd^ (O3) and 4^th^ (O4) harmonics show the absence, independently from the depth, of amide bands in the protein-free region. On the contrary, in the ellipse-circled area, amide signals decrease going deeper into the fibril, as indicated by the diminishing amide/DTTO ratio.

Depth profiling via higher harmonic nano-FTIR spectra (2^nd^, 3^rd^, and 4^th^ harmonics) was performed to provide volumetric chemical information (Fig. 4E-F) ^[26]^. The second harmonic demodulation of the cantilever frequency penetrates deepest into the sample, the third harmonic somewhat less, and the fourth harmonic is the most surface sensitive. The averaged spectrum across the flat zone of the fibril (Fig. 4E) has shown no signs of amide bands in its second to fourth harmonic and therefore proves the flat region to be protein-free at any depth. Conversely, the rough region has shown strong amide signals confined to the fibril surface, with the amide-to-DTTO peak ratio decreasing with increasing depth (Fig. 4F, green spectra: 4^th^ to 2^nd^ harmonic), indicating a proteinaceous outer layer surrounding a pure DTTO-rich core.

### DTTO fibril serve as conductive interface to modulate neuronal function

To assess whether the presence of the fibrils could alter neuronal electrical properties, whole-cell patch clamp recordings were conducted on undifferentiated SH-SY5Y cells containing either monomeric DTTO or assembled fibrils (Fig. 5A-C). Cells with DTTO fibrils embedded exhibited a marked increase in membrane capacitance (Cₘ) (Fig. 5D, Fig S23A), likely arising from an electrostatic coupling between the fibrils and the plasma membrane, and a steeper input resistance slope (RInput) (Fig. 5E, Fig. S23B). Notably, cells with fibrils also resulted in a depolarized resting membrane potential (RPM) (Fig. 5F, Table S1), which is best explained as a downstream consequence of higher capacitance rather than a direct alteration of ionic gradients. An increased Cₘ is expected to prolong the membrane’s time constant, thereby facilitating temporal integration of sub-threshold oscillations and causing a shift of the steady-state potential toward more depolarized values.

**Fig. 5.**
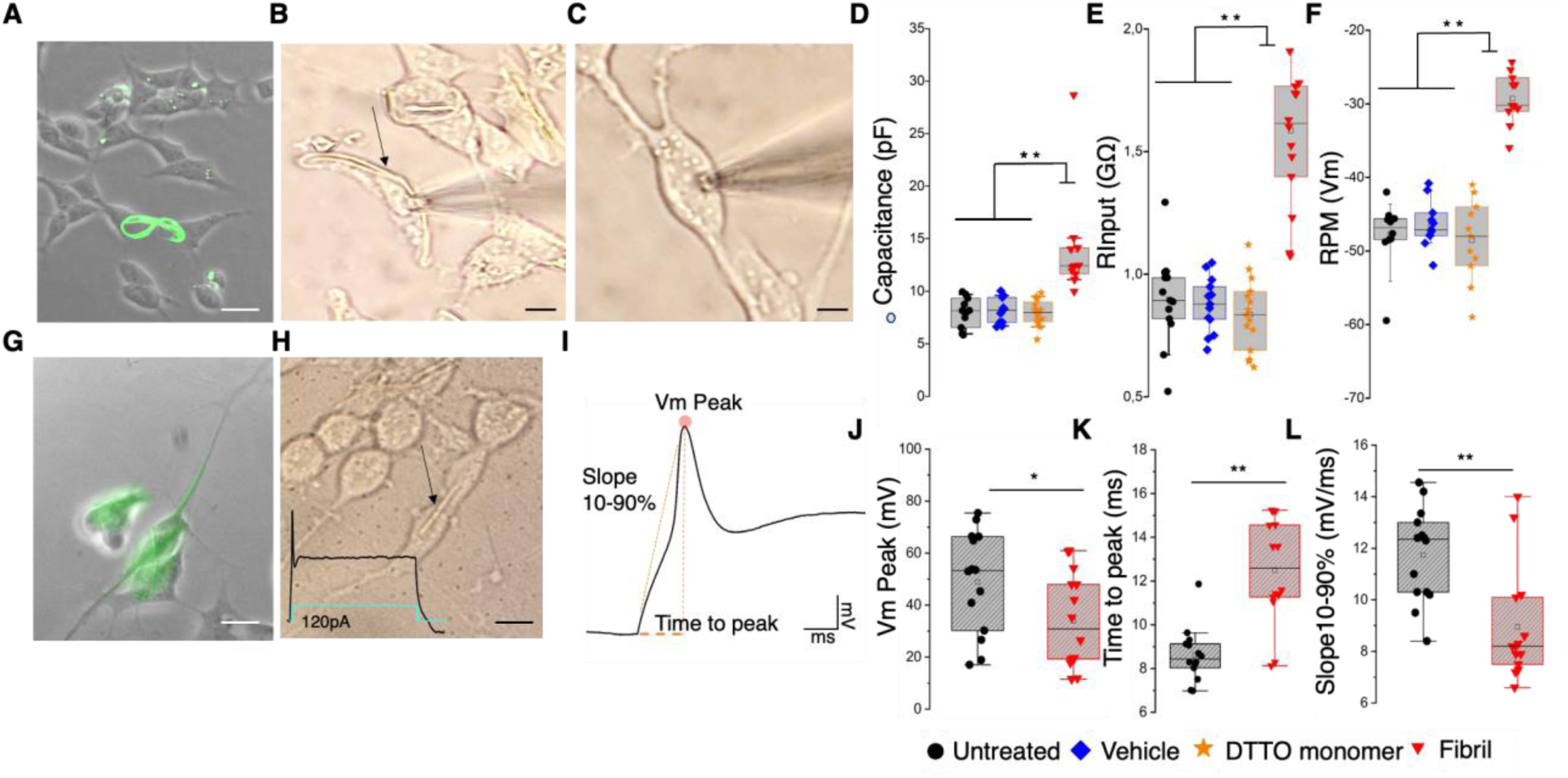
DTTO fibrils modulate neuronal excitability. **(A-C)** Representative undifferentiated SH-SY5Y cells employed for patch clamp recordings: **(A)** fluorescence-brightfield merged image of DTTO treated cells, showing a flake-shaped fluorescent fibril between two cells. Scale bar: 20 µm. **(B)** Brightfield images of cells containing the neo-formed fibrils and **(C)** the DTTO monomer. Scale bars: 2 µm. **(D–F)** Box plots representing membrane capacitance (Cm, **D**), input resistance (RInput, **E**), and resting membrane potential (RPM, **F**). Numerical values are reported in Table S1. **(G–H)** RA-differentiated SH-SY5Y cells: **(G)** representative fluorescence-brightfield merged image of differentiated neurons showing fibrils extending into the neurites. **(H)** Image of fibril-producing neurons employed for patch clamp recordings. Inset: representative voltage response to suprathreshold current injection (cyan trace). Black arrows indicate the fibril. Scale bars: 20 µm. **(I)** AP parameters analyzed to assess fibril impact. **(J–L)** Box plots depict median (thin horizontal bar) and the 25^th^ and 75^th^ with whiskers showing the 5^th^ and 95^th^ percentile of Vm peak **(J)**, time to peak **(K)** and slope of 10-90% AP rising phase **(L).** One-way ANOVA and Bonferroni’s *post hoc* test, \**p* < 0.05. Values ± SD are reported in Table S2.

In retinoic acid (RA)-differentiated SH-SY5Y cells, exhibiting key neuronal features (Fig. 5G-H, Fig. S24), while passive membrane properties remained unchanged (Table S2, Fig. S23D-F), consistent with established effects of RA-induced differentiation, significant differences emerged in various aspects of the evoked action potential (AP) (Fig. 5I-L). DTTO fibrils specifically modified the early phase of AP initiation (Fig. S23C, G-H), indicating that their primary effect is on passive membrane charging prior to channel opening and, even indirectly, on neuronal excitability. Altogether, these results indicate that DTTO fibrils function as dynamic conductive interfaces, reshaping membrane biophysics and modulating cellular excitability. Such findings underscore the potential of biohybrid materials to influence neuronal computation and network-level responses.

### Conclusions

This study establishes a transformative paradigm in neuroelectronics, showing the fabrication and integration of living electronic interfaces within neurons. By leveraging the cell’s intrinsic machinery, we demonstrate that simple molecular precursors such as DTTO can be autonomously assembled into highly fluorescent microfibrils acting as dynamic interfaces, able to modulate membrane biophysical properties and neuronal excitability. The fibril biogenesis, dissected through a multimodal analysis, is governed by a tightly regulated interplay between lipid droplet sequestration of DTTO monomer and autophagy, resulting in biohybrid architectures tailored by the local cellular milieu. A proposed model arising from our findings is showed in Fig. 6.

**Fig. 6.**
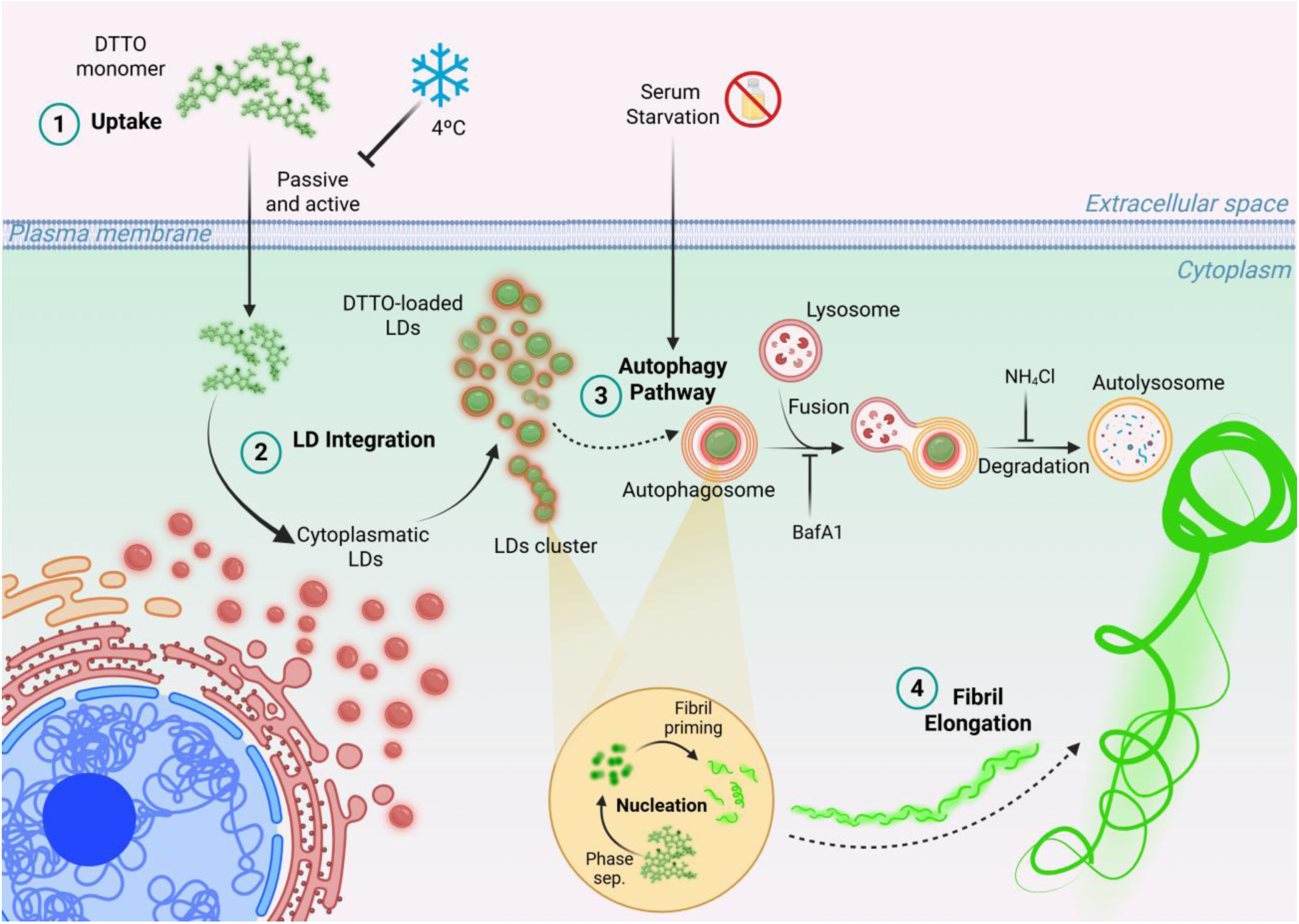
Proposed model for DTTO fibril biogenesis within cells. When interfacing cells, DTTO monomer is internalized *via* passive and active mechanisms (1), as demonstrated by the reduced uptake rate observed under cold temperature incubation. Once inside the cytoplasm, DTTO monomer is directed to the lipid droplets (LDs) pathway (2), being concentrated within the core of cytoplasmatic LDs. The highly hydrophobic environment of LDs core facilitates the local confinement of DTTO monomer up to a critical value which possibly promote the fibril priming. Indeed, LDs may act as nucleation points, favouring fibril initiation through their coalescence and being regulated by lipid autophagy. Limiting LDs degradation and promoting their accumulation (3) via autophagy modulation (serum deprivation, bafilomycin and NH_4_Cl) strongly enhance the fibril yield, acting on their biogenesis. The fibril elongation and supramolecular organization (4) may occur through additional monomer recruiting and might involve intermediate filaments and microtubules.

The presence of proteins on the surface and sub-surface of DTTO fibrils and their exceptional purity and crystalline order are the central features distinguishing these structures as true biohybrid materials rather than simple molecular aggregates. Multiple ultrastructural and spectroscopic evidence support this phenomenon: i) TEM-EDS analyses consistently detect both the sulphur characteristic of the DTTO backbone and the nitrogen, abundant in proteins, within the fibril and especially at helical pitches and at the edges of the fibril structure, indicative of protein incorporation or tight surface association. These spatial patterns are further confirmed by ii) EELS, showing that the nitrogen accumulation is not an artefact caused by sample thickness or technical limitations. Further, iii) nano-FTIR spectroscopy unambiguously identifies the presence of proteins at the surface, detecting amide I and amide II infrared absorption bands in fibrils derived from living cells and not in DTTO aggregates. Clarifying whether the fibril biogenesis is cell type-specific or linked to distinct physiological states will provide deeper insight on how the fibrils influence cell functions and may be used for tailored applications. Here, we exploit their potential in neurotechnology for non-invasive, cell-autonomous modulation of neuronal activity.

Electrophysiological experiments show that cells containing DTTO fibrils acquire distinct electrical signatures, such as a significant increase in membrane capacitance and a more depolarized resting membrane potential compared to untreated cells. The presence of fibrils also modified the early rising phase of the action potential, indicating that DTTO fibrils function as modulatory interfaces possibly due to their conductive properties^[16b, 16c, 17]^. By reshaping membrane electrical properties, they could enhance cellular responsiveness and tune the excitability of neural cells in a controllable fashion. The detection of the fibrils frequently joining adjacent cells, directly through neurites (Fig. 5G, Fig. S24) or creating extracellular bridges (Figs.1A, C; 5A), suggest their capability to generate *in situ* conductive pathways and microcircuits between neurons, writing novel connections into living brains.

Collectively, these findings unlock new strategies for engineering living tissues with built-in electronic functionality, providing a versatile platform not only for neurotechnology, but also for regenerative medicine and synthetic biology. Looking ahead, deciphering and harnessing the principles of biologically directed self-assembly could enable the design of customizable biohybrid devices capable of interfacing with complex systems, toward fully integrated “cyborg” tissues and organs.

## MATERIALS AND METHODS

### Cell culture

Human neuroblastoma SH-SY5Y were purchased by ATCC^®^ (CRL-2266), cultured in Dulbecco’s modified Eagle’s medium (DMEM - Gibco™ 12491015) supplemented with 10% of fetal bovine serum (FBS) (Gibco™ 11560636), 5% of GlutaMAX™ (Thermo Scientific™ 35050038) and 5% of penicillin/streptomycin antibiotics (Gibco™-15070063) and maintained in a humidifier incubator at 37 °C in a 5% CO_2_ atmosphere.

### DTTO cell treatment

DTTO oligothiophene used in this work has been developed and synthetized starting from commercial precursors as previously reported ^[15]^. The DTTO emission spectrum exhibits a main peak at approximately 408 nm, while the emission spectrum shows a maximum at 514 nm. DTTO was solubilized in dimethyl sulfoxide (DMSO) to a final concentration of 1 mg/mL. SH-SY5Y neuroblastoma cells were seeded in multiwell plates or on Poly-D-Lysine (PDL) coated glass coverslips and cultured 48 hours before treatment with DTTO at concentration ranging from 5 μg/mL to 50 μg/mL in DMEM without FBS, unless otherwise specified. After treatment, cells were extensively washed with DPBS 1X (Gibco™-14040133), maintained in DMEM supplemented with FBS and monitored at different times according to the experiment.

### *In vitro* cytotoxicity assay

SH-SY5Y cells were seeded in 96-well plates (1 × 10^4^ cells/well) and cultured for 48 hours. Cells were then treated with DTTO at various concentrations (5, 25 and 50 μg/mL) for 30 min, 1 hour and 16 hours (only at 5 μg/mL) at 37 °C and 5% of CO_2_. Untreated and vehicle-treated cells with equivalent concentration of DMSO (5, 2.5 and 0.5% v/v) were used as controls. Cells were thoroughly washed with DPBS and MTT assay was performed immediately after treatment or 24 hours post treatment (only at 50 μg/mL). MTT (3-(4,5-Dimethylthiazol-2-yl)-2,5-Diphenyltetrazolium Bromide) solution (ThermoFisher M6494) at a final concentration of 0.5 mg/mL in DMEM w/o phenol red was added to each well. Following 60 min of incubation at 37°C, plates were centrifugated (1250 rpm for 20 min) and formazan salts were dissolved with 100 μl of DMSO in agitation for 15 min; the resulting absorbance was determined at λ = 560 nm on a microplate reader (GloMax® Discover Microplate Reader, Promega). Each condition was analyzed in quintuplets and data are expressed as the average percentage of cell viability normalized for the untreated control condition.

### Flow cytometry

Following 1-hour treatments (5, 25 and 50 μg/mL), cells were detached by trypsinization, collected by centrifugation at 350 g for 6 min and resuspended in PBS 1X. A flow cytometer (CytoFLEX V0-B4-R0 Flow Cytometer, Beckman Coulter) equipped with an active laser (488 nm) and 4 detection channels was employed. A total of 1×10^4^ events of single cells were collected for the analysis. FCS files were analysed in FlowJo (BD Biosciences). To eliminate debris, cells were first gated as forward scatter-area (FSC-A) and side scatter-area (SSC-A), while doublets were excluded using FSC-A vs FSC-H (height) plot. Finally, to evaluate DTTO internalization, FITC signal (FITC-A) was plotted into a histogram, and cells were gated to identify positive and negative stained population. Median fluorescence intensity (MFI) for each condition was taken into account and the MFI ratio for each population was calculated as following:

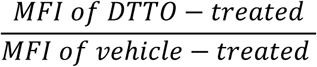

### Holotomography Imaging

Holotomography imaging was performed using a Tomocube HT-2H microscope (Tomocube Inc., Daejeon, South Korea), a digital holographic microscope that enables label-free, quantitative and high-resolution three-dimensional reconstruction of cellular refractive index (RI) distributions (lateral resolution: 110 nm; axial resolution: 220 nm). The HT-2H system integrates three LEDs (wavelength center = 385 nm, 470nm, 570nm) for triple channel fluorescence and a coherent diode laser light source (wavelength: 532nm, output power < 0.4mW) for holographic acquisition. Tomographic RI volumes were reconstructed using a multiple illumination angle via a digital micro-mirror device. Images were acquired using a 60× water-immersion objective (NA 1.2) optimized for live-cell RI measurements, coupled to a high-sensitivity CMOS camera (2048 × 2048 px).

For fluorescence-assisted correlative imaging, Bodipy-labeled lipid droplets were visualized through the red fluorescence channel (excitation 570 nm), while the intrinsic fluorescence of DTTO was detected in the blue-green channel (excitation 470 nm). Fluorescence acquisition was always performed prior to holo-tomographic imaging to minimize photobleaching and to preserve the integrity of RI tomography.

For real-time visualization of fibril biogenesis, SH-SY5Y cells were incubated with 5 μg/ml DTTO, and RI tomograms were acquired every 40 min over the entire imaging period. A stage-top environmental chamber maintained controlled temperature (37 °C) and CO₂ concentration (5%), ensuring physiological conditions throughout prolonged live-cell acquisition. Exposure time for holographic frames was typically 0.3–1.0 ms per illumination angle, and a complete 3D RI tomogram was reconstructed from ∼49 illumination angles in approximately 1–2 seconds, enabling minimally phototoxic time-lapse imaging.

### Holotomography Data Analysis

Image acquisition and 3D reconstruction were performed using TomoStudio software suite (Tomocube Inc.), which controls fluorescence imaging and holo-tomogram acquisition. TomoStudio was used to generate the three-dimensional RI tomograms and to register fluorescence stacks with the corresponding RI volumes through automated correction of chromatic shifts and stage drift.

Quantitative analysis was carried out in TomoAnalysis (Tomocube Inc.), the dedicated post-processing software designed for voxel-level segmentation and numerical extraction of structural parameters. TomoAnalysis enabled measurement of the fluorescent volumes associated with Bodipy-labeled lipid droplets (red channel) and DTTO fluorescence (green channel), as well as their spatial overlap, morphological descriptors, and colocalization with RI-derived structures.

### Immunofluorescence analysis

SH-SY5Y cells were seeded on 13 mm PDL glass coverslips and cultured at least 36 hours. Cells were then treated with DTTO or DMSO using different doses and time, depending on the immunofluorescence assay. After extensive washing with DPBS 1X, cells were fixed with PFA 4% or PFA 2% (tubulins) for 15 min at RT. Following extensive washing in PBS, permeabilization was carried out in PBS/saponin 0.1% (clathrin and perilipin2), PBS/Tween20 0.1% (vimentin) or PBS/Triton X100 0.1% for 15 min. Non-specific bonds were masked by incubation in 3% BSA in PBS/Tween20 0.1% buffer for 1 hour and then cells were incubated with primary antibodies diluted in the same buffer or 1% BSA in PBS/saponin 0.05% (clathrin and perilipin2) for 1 hour at 37 °C. Cells were washed in PBS 1X and then stained with secondary fluorescent antibody diluted 1:200 in the same buffer and incubated 30 min at RT. Finally, after washing the excess of secondary antibody, nuclei were stained with DAPI (1 µg/mL) for 10 min, coverslips were washed in H_2_O and mounted with ProLong™ (Invitrogen P36961).

#### Primary antibody

Monoclonal Anti-α-Tubulin (T6199 Sigma-Aldrich) 1:1000, Monoclonal Anti-β-Tubulin (T4026 Sigma-Aldrich) 1:1000, Anti-clathrin Heavy Chain Antibody, Mouse monoclonal (C1860 Sigma-Aldrich) 1:200, Perilipin-2 (PLIN2) Polyclonal antibody (15294-1-AP Proteintech) 1:400, Monoclonal Anti-Vimentin (V6630 Sigma-Aldrich) 1:1000. Secondary antibody: Goat anti-mouse IgG (H+L) Alexa Fluor™ 555 (A21424 Invitrogen), Goat anti-rabbit IgG (H+L) Alexa Fluor™ 555 (A32732 Invitrogen).

#### CellMask

Cells were stained with CellMask^TM^ Orange Plasma Membrane Marker (2 ug/ml, Invitrogen C10045) for 7 min at 37 °C. After extensive washing with PBS 1X, cells were fixed with PFA 2% and subsequent procedures were performed according to specific aims of the experiment.

#### Lysotracker

Cells were co-treated with 50 nM Lysotracker Red (L7528 Invitrogen) and 5 mg/ml of DTTO at 37 °C for 30 min, protected from light. After incubation, cells were washed once with PBS to remove excess dye, fixed with PFA 4% and processed as above described.

#### Phalloidin staining

Cells were stained with rhodamine phalloidin (R415 Invitrogen) following the manufacturer instruction. Cells were fixed (PFA 4% for 15 min) and permeabilized with PBS/Triton X-100 0.1% for 15 min. After washing with BSA 1% in PBS, cells were stained for 1 hour at RT with phalloidin diluted 1:400 in the washing buffer. Finally, nuclei were stained, and coverslips mounted as previously explained.

#### Bodipy staining

Cells were treated with 1:1000 dilution of Bodipy 558/568 (D3835 Invitrogen) C12 for 24 hours and then with 5 mg/ml of DTTO or a 4-hour coincubation treatment was performed.

### Cell treatments for fibril inhibition: cold shock, starvation and autophagy inhibitors

Cold shock *-* Treatments were carried out either at 4°C or 37 °C for 30 min using DTTO 50 μg/mL. After the necessary time, cells were extensively washed with DPBS 1X to eliminate the DTTO excess. Groups of cells treated at 37 °C were cold shocked for 30 min at 4 °C to inhibit the metabolism and then cultured at 37 °C for all the conditions analysed.

Nutrient starvation *-* To change the nutrient availability, cells were either maintained in FBS-supplemented media or kept in FBS-deprived media for 24 hours prior treatment. After that, in both experimental groups, DTTO treatments (5 μg/mL) were carried out either in FBS-supplemented DMEM or serum-free DMEM for 24 hours.

Autophagy inhibition by Bafilomycin A1 (Sigma-Aldrich, B1793) *-* To inhibit autophagy, cells were pre-incubated with BafA1 100 nM in DMEM with FBS for 30 min. DTTO treatments (5 μg/mL) in presence of BafA1 in DMEM without FBS were performed for 2 hours.

Autophagy inhibition by ammonium chloride (NH_4_Cl) *-* Cells were pre-incubated with NH_4_Cl 25 mM in DMEM with FBS for 30 min and then treated with DTTO 50 μg/mL in DMEM without FBS for another 30 min.

### Quantification analysis

Evaluation of yield fibril production - To calculate the yield, images of 4 areas/well were taken by fluorescence microscopy (Axiovert 100, Zeiss, JENA, Germany) using the software Leica LasX v.3.7.6. The total number of nuclei and fibrils in each area imaged were quantified using Fiji software and the percentage of cells producing fibrils, namely the yield of fibrils synthesis, was calculated using the following equation:

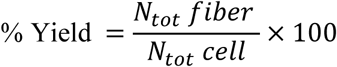

Data represent the average percentage yield values obtained by triplicates for each condition analysed from two independent biological experiments.

Quantification of DTTO uptake - Fluorescence images were taken by the software Nikon NIS-Element using the same exposure, lamp intensity and objective parameters to allow the comparison between various conditions. Different areas were acquired for each condition. The pixel intensities curves were obtained from selected ROI using Fiji software. The list of grey values and the number of pixels for each value were plotted in an XY distribution and the curves for each condition analysed were fitted by means of Gaussian function using Prism 9 (GraphPad Software Version 9.5.0). To estimate the DTTO uptake rate, the average fluorescence intensity was calculated for each condition. First, fluorescence intensities were quantified from identical ROI per condition (two or three different areas/condition) using Fiji. Mean intensity values obtained were finally corrected for the background mean value of each image. Quantification of fibril length and area - Fibril length (Lf) and area (Af) were measured by Fiji software using the segmented line tool for the length or by applying a mask to specifically recognise each fibril in the image to estimate the area.

Quantification of lipid droplets number, size and DTTO loading - To evaluate the percentage of PLIN2 colocalization with DTTO granules, the total number of LDs and their size, CellProfiler™ v.5 software was used for the analysis of fluorescent images obtained by PLIN2 immunofluorescence assay. Colocalization measurement ^[27]^: to calculate the amount of DTTO monomer stored inside LDs, the degree of colocalization between PLIN2 and DTTO signal was measured by means of Pearson’s correlation coefficient (PCC). Fluorescent images were analysed with a custom-made pipeline adapted by CellProfiler software to identify the total number of LDs and DTTO granules in the whole area screened. PCC was calculated for each identified object and values > 0 were considered as positive colocalization correlation, then identifying the number of DTTO monomer internalized within LDs (PLIN2+DTTO). Finally, the percentage of DTTO within LDs has been calculated as following:

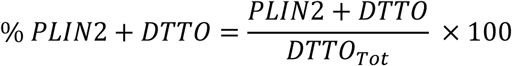

Quantification of LDs area and diameter: the LD area and diameter was calculated by means of a custom-made CellProfiler pipeline. First, all the LDs and nuclei in the screened areas were identified to obtain the total number of LDs and cells. The Area and diameter were measured for all LDs identified and object with eccentricity values between 0 > e < 0.8 were filtered out and not taken into account for the analysis.

Colocalization analysis of DTTO fibrils and cytoskeleton proteins - To evaluate the correlation between the different cytoskeletal markers studied and the DTTO fibrils, two different colocalization analysis were performed:

Object-based colocalization analysis: correlation between fibrils and the cytoskeletal markers was firstly evaluated on the whole population of cells observed in the image by means of an object-based colocalization approach. The degree of correlation was assessed by calculating the PCC for each fibril by means of a custom-made pipeline developed in CellProfiler™ v.5 software. Fibrils associated to saturated signals were excluded from the analysis while the number of colocalizing fibrils was obtained considering values of PCC > 0.1. The percentage of Colocalized/Not Colocalized fibrils was calculated using the following equation:

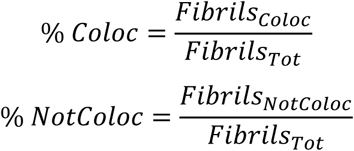

Pixel-wise colocalization analysis: to evaluate the correlation on a single representative fibril, a manual pixel-wise analysis was performed on a line ROI. Intensity pixel values along the ROI were extracted from both the GFP and TRITC channels using a Fiji Macro available on <u>GitHub</u> (Macro_plot_lineprofile_multicolor Kees Straatman, University of Leicester, Leicester, UK). Intensity values from both channels were then plotted against each other in a scatter plot. Correlation analysis and linear regression were finally performed using Graphpad Prism software (version 9.5 for Mac) to calculate PCC and R^2^ values.

Colocalization analysis of confocal and super-resolution imaging – Imaging was carried out using a Nikon Ti2 inverted microscope (Nikon Europe B.V, Amsterdam, The Netherlands) equipped with the CrestOptics X-Light V3 spinning disk system, the DeepSIM X-Light super-resolution add-on module (CrestOptics, Rome, Italy), the Celesta multi-mode multi-line laser source (Lumencor, Beaverton, OR, USA), and the Kinetix sCMOS camera (Teledyne Photometrics, Tucson, AZ, USA). Colocalization analysis to evaluate correlation between DTTO fibrils and tubulins’ signal was performed using JACoP v2.0 (Just Another Colocalization Plugin)^3^ plugin for Fiji software on maximum projections of Z-stack planes and cytofluorograms and Mander’s coefficient of representative fibrils were obtained. Cytofluorogram values were then processed in GraphPad to calculate with correlation analysis the relative Pearson’s coefficient. 3D reconstructions were obtained using Nikon NIS-element software.

### DTTO fibrils isolation from cell cultures

Fibrils were extracted from SH-SY5Y cells cultured in T25 flask and treated with DTTO overnight (5 µg/mL). After treatment, culture was washed twice with cold PBS 1X and cells were collected with a scraper and resuspend them in 2 mL of hypotonic buffer for cell lysis (10 mM HEPES, 10 mM KCl, 1.5 mM MgCl_2_, 0.05 % Tween 20, pH 7.9) containing protease inhibitor cocktail (P8340 Sigma-Aldrich). Cells were left on ice for 20 min and then cell’s lysate was collected and centrifuged for 10 min at 3000 rpm at 4 °C to separate cytoplasmatic from nuclei fraction. Supernatant containing cytoplasmatic fraction was collected and further centrifuged in the same way to eliminate possible debris or remaining nuclei. Pellet was incubated in the same hypotonic buffer while supernatant containing fibrils was collected. As fibrils quantity may be too low, to concentrate them, all supernatants (*i.e.* also the one derived from the second pellet) were centrifuged together at 5000 rpm at 4 °C. To follow isolation steps, fibrils presence after each passage was assessed by fluorescence microscopy. During the last steps, fibrils concentrated in pellets were washed and resuspended in H_2_O.

### Morphological and elemental analysis by Cryo-TEM, TEM-EDS and TEM-EELS

Fibrils isolated by hypotonic lysis were analysed by Cryo-TEM, TEM-ESD and TEM-EELS.

Cryo-TEM *–* 5 µL of fibril suspension was applied to glow-discharged Quantifoil® R 1.2/1.3 300 Mesh Cu grids (Quantifoil, Großlöbichau, Germany) and was flash frozen in liquid ethane using a Vitrobot mark IV (Thermo Fisher Scientific, Waltham, Massachusetts, USA) set at 100% humidity and 25° C for the preparation chamber 2 s blot time. Cryo-EM micrographs were acquired on a Glacios microscope (Thermo Fisher Scientific) operated at an accelerating voltage of 200 kV with a 50 µm C2 aperture, at an indicated magnification of 11.5k, 39k and 69k X in nanoprobe EFTEM mode with a total dose of 20 and 50 e/Å2 and a defocus value range of −3 µm and −5 µm. The direct electron detector was a Falcon 4i positioned post a Selectris energy filter (Thermo Fisher Scientific), operated in a zero-energy-loss mode with a slit width of 10 eV. The beam was set up in parallel illumination by diffraction pattern on carbon foil to reduce artifacts. Before acquisition astigmatism and coma were check by using Sherpa software and the energy filter was aligned to zero loss point for each analyzed grid square. Images were acquired by using Smart EPU software (Thermo Fisher Scientific), the selected images were analyzed by Velox software (Thermo Fisher Scientific).

TEM-EDS and TEM-EELS *–*10 or 15 µL of fibrils suspension were spotted on TEM Holey carbon-coated grid and air-dried. TEM-EDS/EELS analysis was performed (Thermo Fisher Scientific) in the ELECMI facility at the DME (Division of Electron Microscopy) located in the University of Cadiz (ES). Talos F200X G2 TEM/STEM, equipped with a high brightness field emission gun (XFEG), high efficiency XEDS system (Super-X G2) and 4 different STEM detector [HAADF, MAADF (DF4), ADF (DF2) and BF] was used for TEM-EDS analysis. TEM/STEM FEI Cubed Titan Themis, equipped with a high brightness field emission gun (XFEG), direct electron detection gatan K3-IS system was utilized for TEM-EELS. Images were acquired by high sensitivity Ceta CMOS camera and Velox software was used for elemental analysis (Thermo Fisher Scientific). For both TEM-EDS and TEM-EELS, merged images were analysed by Fiji software and profile plots from different elements were obtained using the Macro available on <u>GitHub</u> (Macro_plot_lineprofile_multicolor Kees Straatman, University of Leicester, Leicester, UK).

### Nano-FTIR spectroscopy

For nano-FTIR characterization, 20 µL of fibrils suspension and 5 µL of DTTO monomer (1 mg/mL) suspension were spotted on silica wafers and air-dried overnight. Silica wafers were then washed rapidly with water, dried and analysed by nanoscale IR-spectroscopy with the *IR*-neaSCOPE^+s^ system by neaSPEC (attocube system GmbH)^[27–28]^. Pt−Ir coated AFM tips (typically 275 kHz) with an apex radius of about 25 nm were used with a tapping amplitude of about 70−90 nm. As light source a tunable broadband nano-FTIR laser (based on difference frequency generation) ranging from 650 cm^-1^ to 2200 cm^-1^ served for mid-IR illumination. In the nano-tomography analysis peak height ratios between the amide I band at 1656 cm^-1^ and the DTTO absorption at 1302 cm^-1^ in the nano-FTIR (imaginary part of the dielectric function) spectra at different harmonics were calculated^[29]^ and demonstrated that the protein layer is on top on the otherwise pure DTTO fibers. Forming the peak height ratios in the optical near-field phase spectra arrive at the same result.

### Differentiation of SH-SY5Y neuroblastoma cells

SH-SY5Y cells were differentiated using previously reported protocols^[30]^. Briefly, cells were seeded onto PDL glasses at the specified densities relevant to the experiment. After 48 hours, cells were incubated in DMEM supplemented with 3% FBS and 10 μM retinoic acid (RA) (R2625 Sigma-Aldrich) for 14 days. The RA was dissolved in DMSO to a stock concentration of 5 mM and added to the cell culture medium to the final selected concentration.

### Electrophysiology

SH-SY5Y cells were seeded at concentration of 3×10^3^ cells on 8 mm glass coverslips and grown in 12 multiwell plates. Before electrophysiological experiments, cells were incubated either for 30 min or overnight with 5 µg/mL of DTTO.

Patch-clamp setup and solutions - During electrophysiological experiments, cells were visualized by using a Nikon Diaphot 300 Inverted Phase Contrast Microscope equipped with a 40x objective, mounted with a recording chamber. Patch-clamp recordings in whole cell configuration were performed on cells with internalized fibers that did not protrude through the membrane using a HEKA EPC7 patch amplifier, digitized via a Digidata1200A interface, and acquired with pClamp 8 (Axon Instruments). Signals were sampled at 10 kHz, filtered at 3 kHz and analysed using Clampfit10 and OriginPro2022. Borosilicate glass pipettes (Clark Electrical Instruments) were pulled using a P1000 micropipette puller (Sutter Instruments) and heat-polished before use. Whole-cell recording pipettes had an open tip resistance of 3-5 MΩ with the classical intra- and extracellular solutions (intracellular (mM): 130 KCl; 5 NaCl; 1 CaCl_2_; 2 MgCl_2_; 2 Mg_2_-ATP; 0.5 Na_3_-GTP; 10 EGTA; extracellular (mM): 135 NaCl; 5 KCl; 2 CaCl_2_; 1 MgCl_2_; 10 Glucose; 10 HEPES).

Sucrose was added to solutions to adjust milliosmolarity to 304 and 315 mOsm/L respectively for intra and extracellular recording solutions. The pH adjustment at the physiological value of 7.3 was made with NaOH or KOH. The calculated liquid junction potential with these solutions was about −3 mV and the data presented were not corrected.

Experimental protocol and statistical analysis - Capacitance values were determined by applying a voltage-clamp step of −60 mV from a holding potential corresponding to the RPM value without correction for whole cell capacitance and series resistance (Platzer and Zorn-Pauly, 2016). The resulted decaying current was fitted by a single exponential:

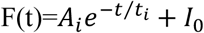

where (*A_i_*) represents the amplitude current, *t* is the time from the onset of the pulse and (*I_0_*) the offset amplitude, to extrapolate the current value (*A_0_*) reached at the beginning of the voltage step (the Y intercept value). Membrane resistance (*R_m_*) at which the peak current (*A_0_*) occurred was calculated by Ohm’s law and finally membrane capacitance (*C_m_*) by the formula:

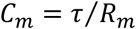

The resting membrane potential (RPM) and the membrane input resistance (R_INPUT_) values were collected in current clamp at holding current of 0 pA. The RPM was registered in gap free mode while R_INPUT_ was estimated as the slope of the linear V-I relationship between the intensity (I) of applied hyperpolarizing currents (15 steps of 100 ms in 5 pA decrements starting from −10 to −80 pA, hc 0 pA) and the corresponding membrane potential measured at each step (V_step_).

Electrophysiological properties were investigated in current-clamp (CC) mode by a 400 ms long current protocol in which a current step of 120 pA was applied at a holding current to set the membrane potential at approximately −100 mV. For each evoked action potential (AP) we quantified the peak membrane potential (Vm peak) defined as the maximum voltage reached during the AP, the latency between the onset of the CC protocol and the occurrence of the peak membrane potential (Time to peak), and the slopes of the rising phase by linear regression over distinct regions (from 10% to 90% of the rising phase, Fig.5H, from baseline to the inflection point, and from the inflection point to the Vm peak).

To ensure recording quality, the following criteria were consistently applied across all experiments: leakage current of the initial seal resistance <0.01 nA at −80 mV, holding currents in whole-cell mode < 50 pA, access resistance not exceedingly twice the initial electrode resistance, and stable RPM.

All data are represented as box and whiskers plots, with the box depicting the mean, thin horizontal bars representing the median (Q2), the 25^th^ (Q1) and 75^th^ (Q3) quartile and whisker showing the 5^th^ and 95^th^ percentile. Alternatively, data are presented as means ± standard deviation (SD).

## Acknowledgments

The authors thank Giuseppe Cacace (Istituto di Scienze Applicate e Sistemi Intelligenti, CNR) for technical assistance. Figure 6 is created with Biorender.com. The authors are also grateful to the Advanced Light Microscopy facility at CEINGE (Napoli, Italy) for providing access to imaging systems and CrestOptics (Rome, Italy) for confocal fluorescence imaging using super-resolution. Authors acknowledge the use of instrumentation as well as the technical advice provided by the National Facility ELECMI ICTS, node Division de Microscopia Electrónica (DME) at Universidad de Cádiz, and Aina Reich (Attocube systems GmbH) for scientific and technical support for nanoFTIR analyses.

## Author contributions

Conceptualization: GT, AT, MM, CT

Methodology: GT, MI, SS, MAF, EE, GC, AT, MM, CT

Investigation: GT, MI, SS, PS, DI, GP, MB, GS, FDM, MZ

Visualization: GT, MI, SS, PS, DI, GP, MB

Funding acquisition MAF, CT

Supervision: MAF, AT, MM, CT

Writing – original draft: GT, SS, PS, CT

Writing – review & editing: GT, MI, SS, DI, MB, MAF, PS, AT, MM, CT

## Supplementary Materials

Supplementary Text Figs. S1 to S24 Tables S1 to S2 Movies S1 to S8

## References

[1] S. Oh, J. Jekal, J. Liu, J. Kim, J. U. Park, T. Y. Lee, K. I. Jang, Adv Funct Mater 2024, 34.

[2] X. Tang, H. Shen, S. Y. Zhao, N. Li, J. Liu, Nat Electron 2023, 6, 109.

[3a] Z. Lin, J. C. Garbern, R. Liu, Q. Li, E. Mancheno Juncosa, H. L. T. Elwell, M. Sokol, J. Aoyama, U. S. Deumer, E. Hsiao, H. Sheng, R. T. Lee, J. Liu, Sci Adv 2023, 9, eade8513

[3b] Q. Li, R. Liu, Z. Lin, X. Zhang, I. G. Silva, S. D. Pollock, J. R. Alvarez-Dominguez, J. Liu, bioRxiv 2024

[3c] H. Sheng, R. Liu, Q. Li, Z. W. Lin, Y. C. He, T. S. Blum, H. Zhao, X. Tang, W. B. Wang, L. S. Jin, Z. L. Wang, E. M. Hsiao, P. Le Floch, H. Shen, A. J. Lee, R. A. Jonas-Closs, J. Briggs, S. Y. Liu, D. Solomon, X. Wang, J. L. Whited, N. S. Lu, J. Liu, Nature 2025.

[4a] X. Yang, T. Zhou, T. J. Zwang, G. S. Hong, Y. L. Zhao, R. D. Viveros, T. M. Fu, T. Gao, C. M. Lieber, Nat Mater 2019, 18, 510

[4b] P. Le Floch, S. Y. Zhao, R. Liu, N. Molinari, E. Medina, H. Shen, Z. L. Wang, J. Kim, H. Sheng, S. Partarrieu, W. B. Wang, C. Sessler, G. G. Zhang, H. Park, X. Gong, A. Spencer, J. H. Lee, T. Y. Ye, X. Tang, X. Wang, K. Bertoldi, N. S. Lu, B. Kozinsky, Z. G. Suo, J. Liu, Nat Nanotechnol 2024, 19.

[5] S. W. Hwang, H. Tao, D. H. Kim, H. Y. Cheng, J. K. Song, E. Rill, M. A. Brenckle, B. Panilaitis, S. M. Won, Y. S. Kim, Y. M. Song, K. J. Yu, A. Ameen, R. Li, Y. W. Su, M. M. Yang, D. L. Kaplan, M. R. Zakin, M. J. Slepian, Y. G. Huang, F. G. Omenetto, J. A. Rogers, Science 2012, 337, 1640.

[6] W. B. Wang, C. D. Sessler, X. Wang, J. Liu, Accounts Chem Res 2024, 57, 2013.

[7] B. D. Paulsen, K. Tybrandt, E. Stavrinidou, J. Rivnay, Nat Mater 2020, 19, 13.

[8a] E. Stavrinidou, R. Gabrielsson, K. P. R. Nilsson, S. K. Singh, J. F. Franco-Gonzalez, A. V. Volkov, M. P. Jonsson, A. Grimoldi, M. Elgland, I. V. Zozoulenko, D. T. Simon, M. Berggren, P Natl Acad Sci USA 2017, 1142807;

[8b] G. Dufil, D. Parker, J. Y. Gerasimov, T. Q. Nguyen, M. Berggren, E. Stavrinidou, J Mater Chem B 2020, 8, 4221.

[9] G. Tommasini, G. Dufil, F. Fardella, X. Strakosas, E. Fergola, T. Abrahamsson, D. Bliman, R. Olsson, M. Berggren, A. Tino, E. Stavrinidou, C. Tortiglione, Bioact Mater 2022, 10, 107.

[10a] G. Tommasini, M. De Simone, S. Santillo, G. Dufil, M. Iencharelli, D. Mantione, E. Stavrinidou, A. Tino, C. Tortiglione, Science Advances 2023, 9;

[10b] G. Tommasini, M. De Simone, M. Blasio, C. Zenna, A. Tino, E. Stavrinidou, S. Santillo, C. Tortiglione, Adv Mater Interfaces 2025.

[11] X. Strakosas, H. Biesmans, T. Abrahamsson, K. Hellman, M. S. Ejneby, M. J. Donahue, P. Ekstrom, F. Ek, M. Savvakis, M. Hjort, D. Bliman, M. Linares, C. Lindholm, E. Stavrinidou, J. Y. Gerasimov, D. T. Simon, R. Olsson, M. Berggren, Science 2023, 379, 795.

[12] F. Ek, T. Abrahamsson, M. Savvakis, S. Bormann, A. H. Mousa, M. A. Shameem, K. Hellman, A. S. Yadav, L. H. Betancourt, P. Ekström, J. Y. Gerasimov, D. T. Simon, G. Marko-Varga, M. Hjort, M. Berggren, X. Strakosas, R. Olsson, Adv Sci 2024, 11.

[13] T. Abrahamsson, F. Ek, R. Cornuéjols, D. Byun, M. Savvakis, C. Bruschi, I. Sahalianov, E. Miglbauer, C. Musumeci, M. J. Donahue, I. Petsagkourakis, M. Gryszel, M. Hjort, J. Y. Gerasimov, G. Baryshnikov, R. Kroon, D. T. Simon, M. Berggren, I. Uguz, R. Olsson, X. Strakosas, Angew Chem Int Edit 2025.

[14a] J. Liu, Y. S. Kim, C. E. Richardson, A. Tom, C. Ramakrishnan, F. Birey, T. Katsumata, S. C. Chen, C. Wang, X. Wang, L. M. Joubert, Y. W. Jiang, H. L. Wang, L. E. Fenno, J. B. H. Tok, S. P. Pasca, K. Shen, Z. A. Bao, K. Deisseroth, Science 2020, 367, 1372;

[14b] C. D. Sessler, Y. M. Zhou, W. B. Wang, N. D. Hartley, Z. Y. Fu, D. Graykowski, M. G. Sheng, X. Wang, J. Liu, Science Advances 2022, 8;

[14c] A. Q. Zhang, K. Y. Loh, C. S. Kadur, L. Michalek, J. Y. Dou, C. Ramakrishnan, Z. A. Bao, K. Deisseroth, Science Advances 2023, 9.

[15] I. Palamà, F. Di Maria, I. Viola, E. Fabiano, G. Gigli, C. Bettini, G. Barbarella, J Am Chem Soc 2011, 133, 17777.

[16a] I. E. Palamà, F. Di Maria, S. D’Amone, G. Barbarella, G. Gigli, J Mater Chem B 2015, 3, 151;

[16b] I. Viola, I. E. Palamà, A. M. L. Coluccia, M. Biasiucci, B. Dozza, E. Lucarelli, F. Di Maria, G. Barbarella, G. Gigli, Integr Biol-Uk 2013, 5, 1057;

[16c] M. Moros, F. Di Maria, P. Dardano, G. Tommasini, H. Castillo-Michel, A. Kovtun, M. Zangoli, M. Blasio, L. De Stefano, A. Tino, G. Barbarella, C. Tortiglione, Iscience 2020, 23.

[17] L. Aloisio, M. Moschetta, A. Boschi, A. G. Fleitas, M. Zangoli, I. Venturino, V. Vurro, A. Magni, R. Mazzaro, V. Morandi, A. Candini, C. D’Andrea, G. M. Paternò, M. Gazzano, G. Lanzani, F. Di Maria, Adv Mater 2023, 35.

[18a] M. A. Ferrara, E. Cavalletti, V. Bianco, L. Miccio, G. Coppola, P. Ferraro, A. Sardo, PLoS One 2025, 20, e0322960;

[18b] K. Kim, W. S. Park, S. Na, S. Kim, T. Kim, W. D. Heo, Y. Park, Biomed Opt Express 2017, 8, 5688.

[19a] A. I. Ivanov, Methods Mol Biol 2008, 440, 15;

[19b] L. D. Cervia, C. C. Chang, L. Wang, F. Yuan, PLoS One 2017, 12, e0171699.

[20a] E. Y. Jeong, H. J. Kim, S. Lee, Y. Park, Y. M. Kim, J Opt Soc Am A Opt Image Sci Vis 2024, 41, C125;

[20b] P. A. Sandoz, C. Tremblay, F. G. van der Goot, M. Frechin, PLoS Biol 2019, 17, e3000553.

[21] T. C. Walther, R. V. Farese, Jr., Annu Rev Biochem 2012, 81, 687.

[22a] R. Singh, S. Kaushik, Y. J. Wang, Y. Q. Xiang, I. Novak, M. Komatsu, K. Tanaka, A. M. Cuervo, M. J. Czaja, Nature 2009, 458, 1131;

[22b] I. Ralhan, C. L. Chang, J. Lippincott-Schwartz, M. S. Ioannou, J Cell Biol 2021, 220.

[23] B. Reynes, E. M. van Schothorst, J. Keijer, A. Palou, P. Oliver, Sci Rep 2019, 9, 19985.

[24] B. Li, C. Sun, J. Sun, M. H. Yang, R. Zuo, C. Liu, W. R. Lan, M. H. Liu, B. Huang, Y. Zhou, Stem Cell Res Ther 2019, 10, 118.

[25] C. Mauvezin, T. P. Neufeld, Autophagy 2015, 11, 1437.

[26a] A. A. Govyadinov, S. Mastel, F. Golmar, A. Chuvilin, P. S. Carney, R. Hillenbrand, ACS Nano 2014, 8, 6911;

[26b] F. Mooshammer, M. A. Huber, F. Sandner, M. Plankl, M. Zizlsperger, R. Huber, ACS Photonics 2020, 7, 344.

[27] B. Knoll, F. Keilmann, Nature 1999, 399, 134.

[28] F. Huth, A. Govyadinov, S. Amarie, W. Nuansing, F. Keilmann, R. Hillenbrand, Nano Lett 2012, 12, 3973.

[29] L. Mester, A. A. Govyadinov, S. Chen, M. Goikoetxea, R. Hillenbrand, Nat Commun 2020, 11, 3359.

[30] R. F. Simoes, R. Ferrao, M. R. Silva, S. L. C. Pinho, L. Ferreira, P. J. Oliveira, T. Cunha-Oliveira, Food Chem Toxicol 2021, 149, 111967.

