## Supplementary Material for "Built-in integrated living electronics: from biosynthesis to modulation of neuronal function"

[1] Istituto di Scienze Applicate e Sistemi Intelligenti "E.Caianiello", CNR; Pozzuoli, Italy.
[2] Instituto de Nanociencia y Materiales de Aragón, INMA CSIC-Universidad de Zaragoza; Zaragoza, Spain.
[3]Attocube systems GmbH; Munich, Germany.
[4] Istituto per la Sintesi Organica e la Fotoreattività, CNR; Bologna, Italy.
[5] Centro de Investigación Biomédica en Red de Bioingeniería, Biomateriales y Nanomedicina (CIBER-BBN); Madrid, Spain.

**This PDF file includes:**
Supplementary Text
Figs. S1 to S24
Tables S1 to S2
Captions for Movies S1 to S8

**Other Supplementary Materials for this manuscript include the following:**

Movies S1 to S8

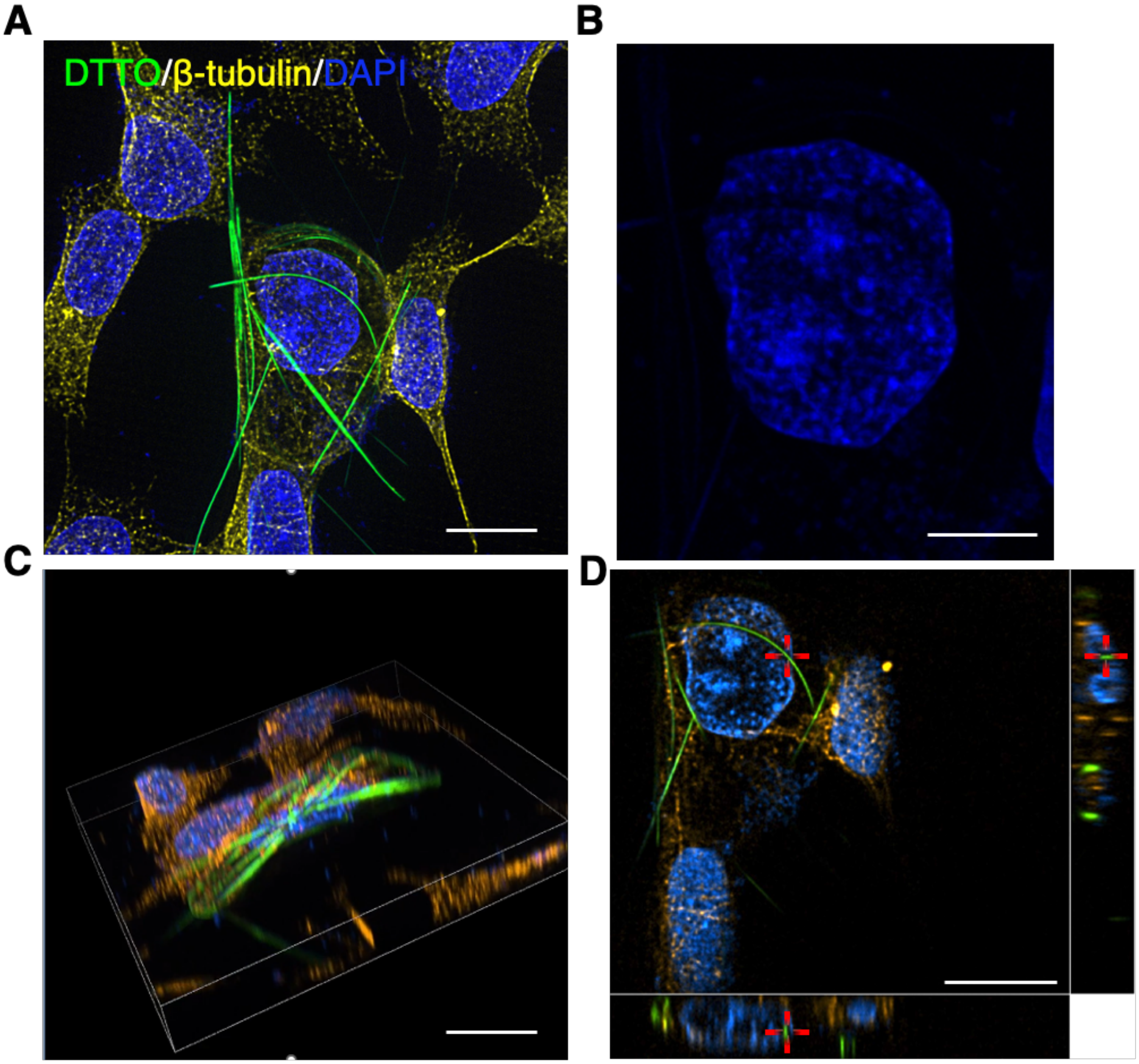


**Fig. S1. DTTO fibrils can penetrate cell nucleus.**

**(A)** Super-resolution image of SH-SY5Y cells incubated overnight with DTTO (5 μg/mL), showing DTTO fibrils (green), β-tubulin (yellow), and nuclei (blue). Scale bar: 10 μm. **(B)** Relative image of the nucleus with a DAPI-free area complementary to the fibril signal, scale bar: 5 μm. **(C)** 3D confocal reconstruction of a cell showing DTTO fibrils (5 μg/mL, overnight) after immunostaining with β-tubulin, scale bar: 10 μm. **(D)** Orthogonal XZ and YZ projections of confocal super-resolution stack showing DTTO and nuclei signals on the same plane, scale bar: 10 μm

**DTTO influence on cell viability, fibril formation, and intracellular uptake: dose and time dependency.**

A systematic quantitative analysis was performed to assess how DTTO monomer dose and incubation time affect cell viability and fibril yield in this specific line. MTT assay on cells treated with DTTO revealed transient cytotoxicity at the highest dose (50 μg/mL), with a 30–50% reduction in viability within 1 hour (Fig. S2), which was primarily attributed to the vehicle (DMSO). However, cells recovered to 90% viability 24 hours post-treatment (p.t.), while no cytotoxic effects were detected following continuous incubation with low dose (5 μg/mL), thereby confirming the absence of long-term negative effects on cell viability from either the monomer or the fibril (Fig. S2D-E). Next, the fibril yield was estimated as function of DTTO dose and incubation time. Higher DTTO dose and longer incubation time increased the fibril yield, with up to 12% of cells producing fibrils following 1-hour incubations, while low dose required longer periods, ultimately yielding up to 24% (Fig. S3). These findings demonstrate that both the intracellular concentration of DTTO and the incubation time are critical for efficient fibril formation. Intracellular DTTO was found both diffusely throughout the cytoplasm and within distinct punctate structures, suggesting sequestration into specific organelles. Fluorescence microscopy and flow cytometry showed that high DTTO concentrations resulted in greater intracellular fluorescence and uptake compared to low doses (Fig. S4A-C). After 1 hour of treatment, over 94% of cells internalized DTTO, with the median fluorescence intensity values increasing proportionally with dose (Fig. S4F). Time-course experiments further revealed that the DTTO uptake rises with the incubation time, especially during the first 20 min (Fig. S5). The concentration-dependent monomer uptake correlates with differences in fibrils synthesis yield, while intriguingly their morphology remained unaffected by DTTO dose, as fibrils produced at 5 μg/ml and 50 μg/ml exhibited comparable dimensions (Fig. S6A). In contrast, significant changes in fibril length and area were observed when increasing the incubation time (Fig. S6B), indicating active processes involved in their elongation over time.

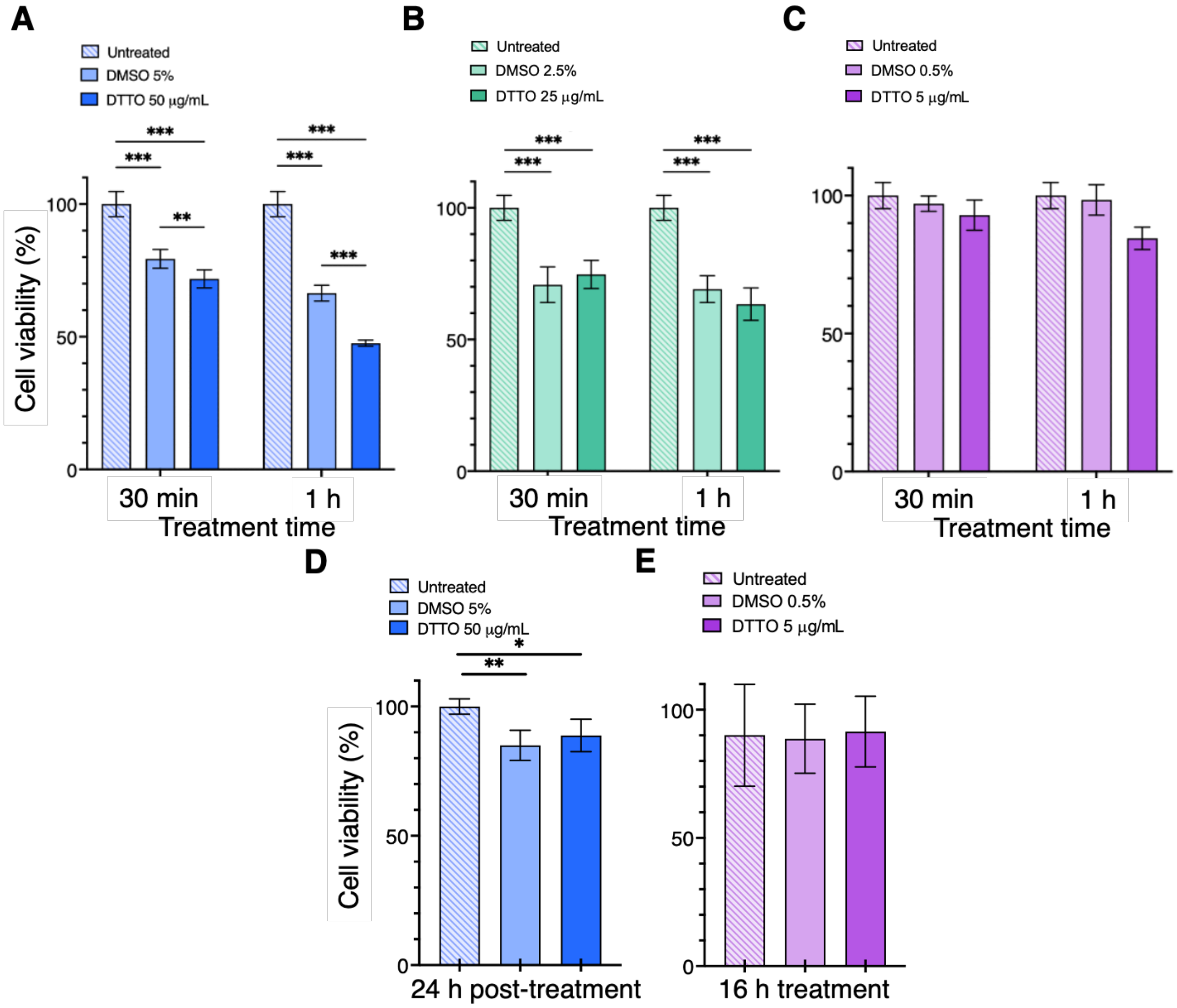


**Fig. S2. Biocompatibility of DTTO as monomer or assembled in fibrils.**

MTT assay performed to evaluate cytotoxicity of DTTO monomer at increasing doses and incubation times. Cells were treated with DTTO at **(A)** 50 μg/mL, **(B)** 25 μg/mL and **(C)** 5 μg/mL for 30 min and 1 hour, and MTT was carried out immediately after treatment. The viability of cells containing embedded fibrils was assessed **(D)** 24 hours p.t. after exposure to DTTO 50 μg/mL for 1 hour or **(E)** after continuous exposure (16 hours) with DTTO 5 μg/mL. Bars represent the mean ± SD of two independent biological experiments, each carried out in quintuplets (n = 10). Mean values for each condition were normalized to untreated control. Statistical analysis was performed using two-way ANOVA followed by Dunnet's post hoc test in **(A**, **B** and **C)** or one-way ANOVA followed by Tukey's post hoc test in **(D** and **E)**; $*p < 0.05$, $**p < 0.01$; $***p < 0.001$.

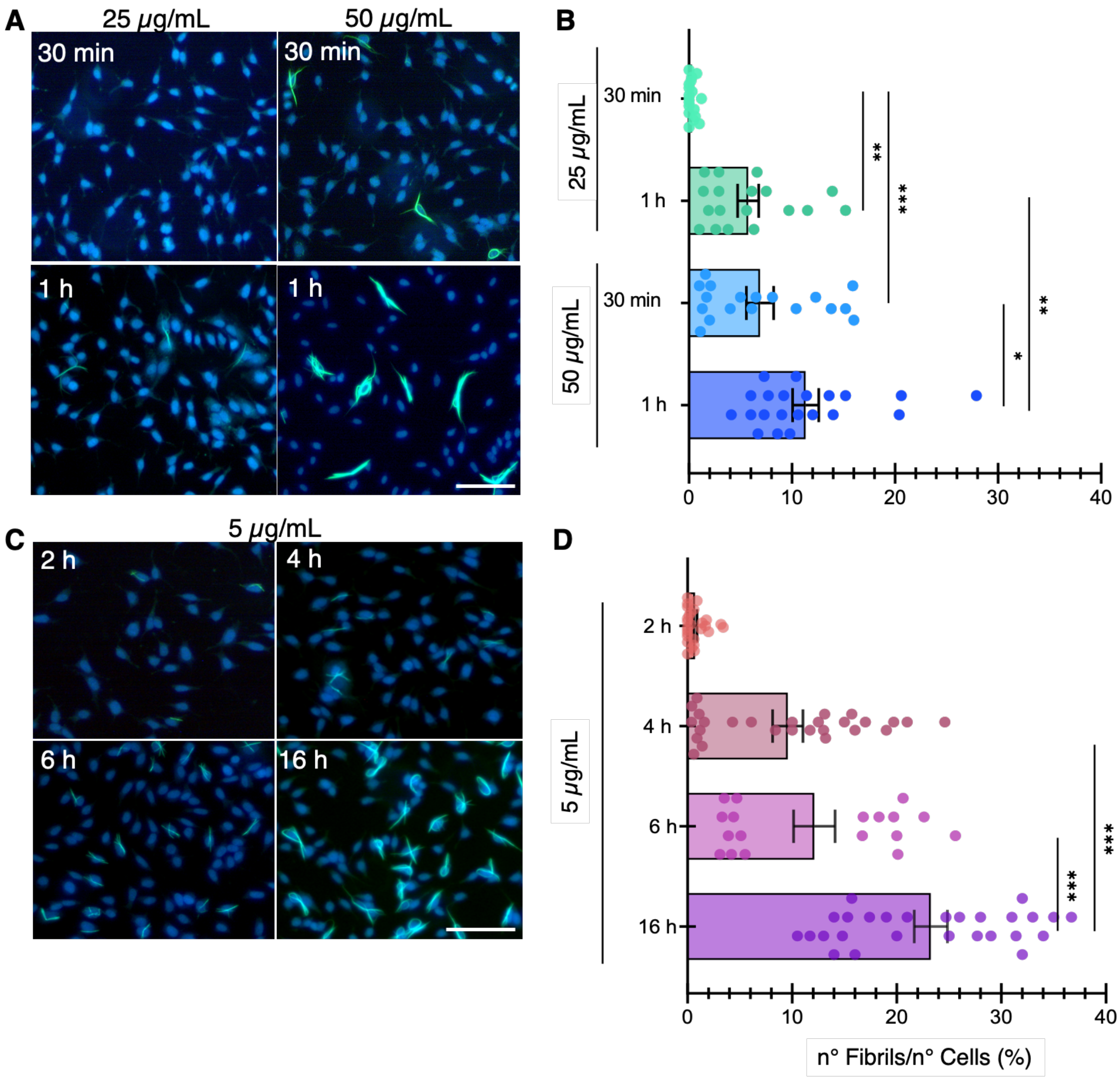


**Fig. S3. Fibril production efficiency is a dose- and time-dependent process.**

**(A)** Representative images of cells treated with 25 and 50 µg/mL of DTTO taken 24 hours post-treatment. After fixation, nuclei were counterstained with DAPI (blue). Scale bars: 100 µm. **(B)** Fibrils were quantified and the percentual yield in each condition was calculated, normalizing the total number of fibrils to the total number of cells (7244 for 50 µg/mL 1 hour, 4373 for 50 µg/mL 30 min, 6876 for 25 µg/mL 1 hour and 6540 for 25 µg/mL 30 min). **(C)** Analysis by fluorescence microscopy showed an increased number of fibrils when cells were treated for extended times up to 16 hours. Images were taken 24 hours post-treatment and nuclei were counterstained with DAPI (blue). Scale bars: 100 µm. **(D)** Fibrils percentage yield after each incubation time was calculated normalizing the total number of fibrils to the total number of cells (5939 for 2 hours, 12733 for 4 hours, 7645 for 6 hours and 10105 for 16 hours). Bars represent the mean ± SEM of two independent biological assays, each repeated in triplicates. Statistical analysis was performed using one-way ANOVA followed by Tukey's post hoc test; $*p < 0.05$; $**p < 0.01$; $***p < 0.001$.

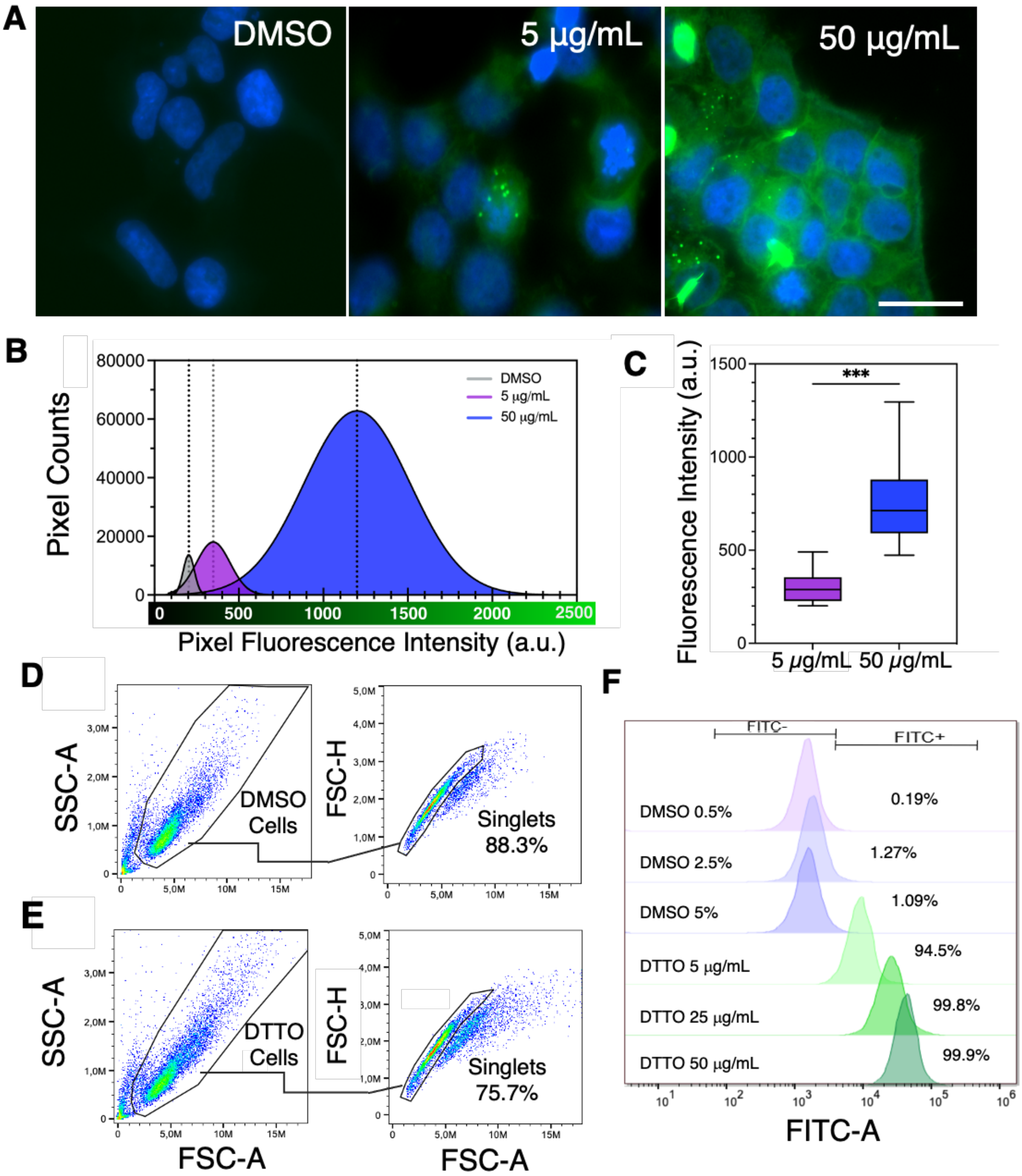


**Fig. S4. Dose-dependent uptake of DTTO monomer**

**(A)** Fluorescence images of cells treated with 5 and 50 µg/mL of DTTO or with vehicle (DMSO) for 1 hour. Cells were fixed and nuclei stained with DAPI (blue). Some short fibrils are already visible within 1 hour of treatment at 50 µg/mL. Scale bar: 25 µm. **(B)** Curves depicting the number of pixels (*i.e.,* count) detected for each intensity value (arbitrary units, a.u.). The data represent the gaussian fitting of the histogram's values from one representative image obtained by Fiji software. Dashed lines represent the mean of the gaussian fitting (DMSO=203.4 a.u.; 5 μg/mL=348.1 a.u.; 50 μg/mL=1199 a.u.). **(C)** DTTO uptake rate at 5 and 50 μg/mL. Data show the fluorescence intensity values calculated from selected ROIs (n=36) within cytoplasm, each representing a single cell, corrected for the background mean; the interquartile range represent the 25$^{th}$ to 75$^{th}$ percentiles while the horizontal solid line represents the median value. Whiskers represent the 5-95% percentiles. Statistical comparison was carried out using Student *t*-test; ***$p < 0.001$. Representative dot plots of **(D)** SH-SY5Y vehicle-treated and **(E)** DTTO-treated (50 μg/mL) cell populations for 1 hour. After treatment, cells were extensively washed and analysed by flow cytometry. **(F)** Dose-dependent internalization of DTTO. Representative histogram overlay of SH-SY5Y cells treated with 5, 25 and 50 μg/mL of DTTO (green shaded curves) for 1 hour. Autofluorescence of vehicle-treated cells (DMSO) is showed as light shaded purples curves. The percentages of DTTO (FITC-A$^{+}$) positive and negative (FITC-A$^{-}$) cells are shown in the graph, while the median fluorescence intensity values are: 0.5% DMSO=1440 a.u.; 2.5% DMSO=1775 a.u; 5% DMSO=1511 a.u; DTTO 5 μg/mL=8729 a.u.; DTTO 25 μg/mL=24764 a.u.; DTTO 50 μg/mL=41044 a.u.

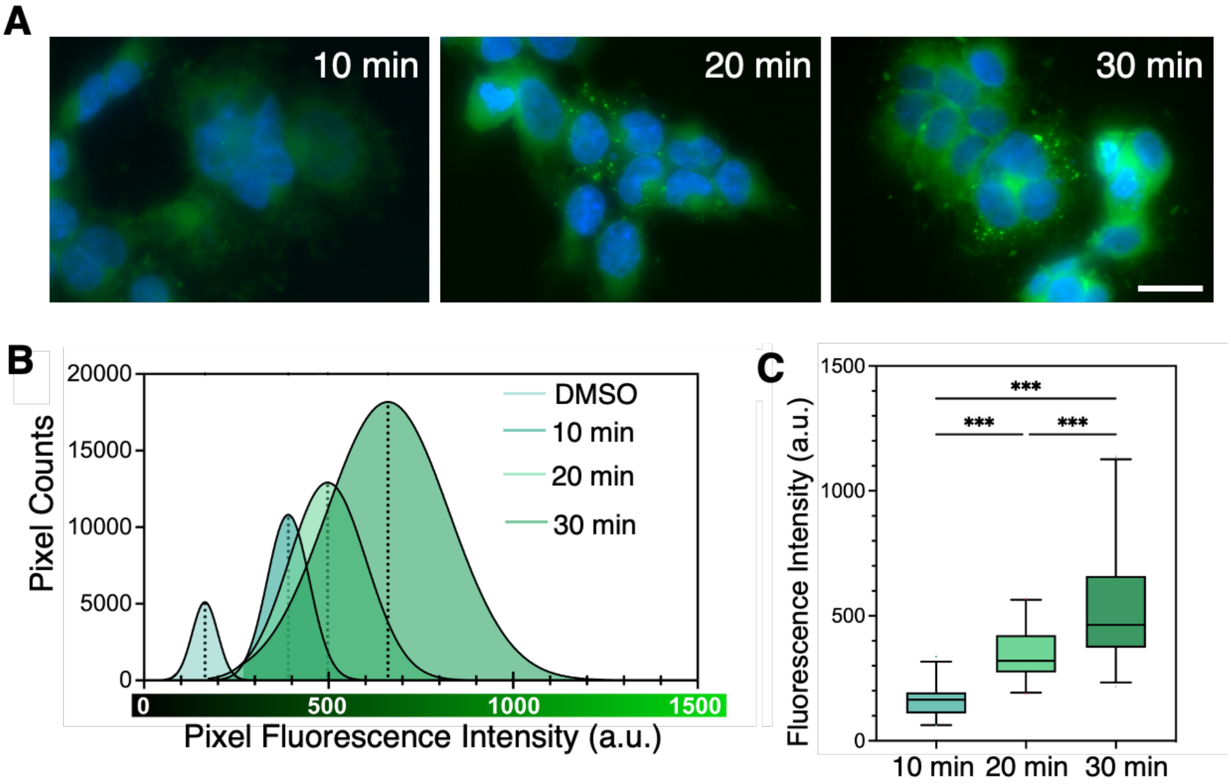


**Fig. S5. Time-dependent uptake of DTTO monomer.**

**(A)** Representative fluorescence images of cells treated with 50 µg/mL of DTTO for 10, 20 or 30 min, fixed, and counterstained with DAPI (blue). Scale bar: 20 µm. **(B)** Fluorescence intensity histograms depicting the number of pixels detected for each intensity value. Curves represent the gaussian fitting of the histogram's values obtained by fluorescence images analysis using Fiji software. Dashed lines represent the mean of the gaussian fitting (DMSO=165 a.u.; 10 min=389.5 a.u.; 20 minutes=497.3 a.u.; 30 minutes=660.7 a.u.). **(C)** Uptake rate at 10, 20 and 30 min. Data show the fluorescence intensity values calculated from selected ROIs (n=25) within cytoplasm, each representing a single cell, and corrected for the background mean; the interquartile range represent the 25th to 75th percentiles while the horizontal line represents the median value. Whiskers represent the 5-95% percentiles. Statistical comparison was carried out using one-way ANOVA followed by Tukey's post hoc test; $***p < 0.001$

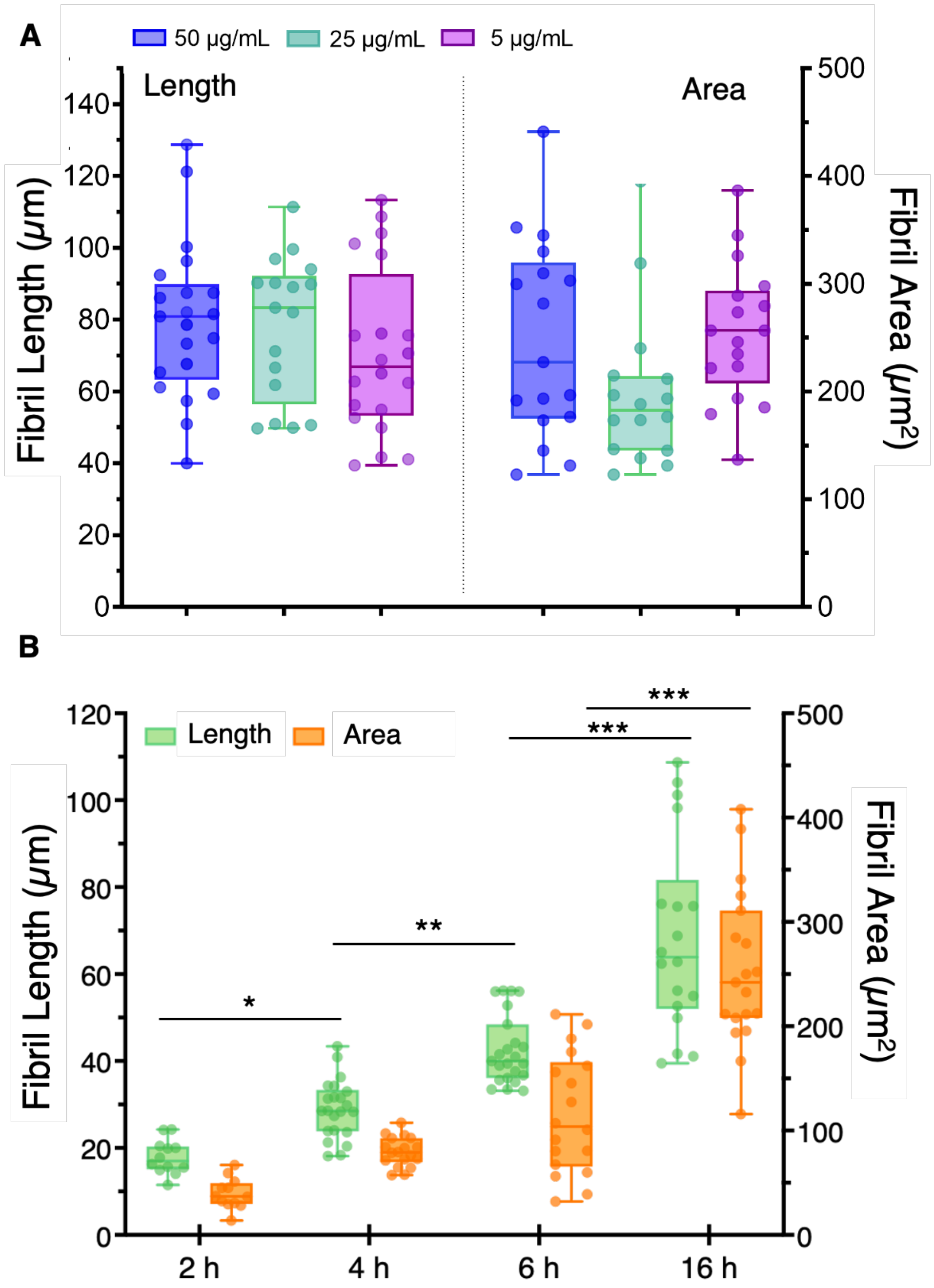


**Fig. S6. Fibrils elongate over time while their size is independent from the dose.**

**(A)** Fibril length and area were calculated at different doses or **(B)** changing the treatment period. The box plots depict the distribution of length and area obtained from two independent biological replicates, while each dot represents a single fibril quantified. Solid line shows the median, while interquartile range represents the 25th to 75th percentiles. Whiskers show the 5-95% percentiles. Length and area were calculated by Fiji software and statistical analysis was performed using one-way ANOVA followed by Tukey's post hoc in (**A**) and Šídák's post hoc test in (**B**); *$p$ <0.05; **$p$ < 0.001; *** $p$ < 0.0001.

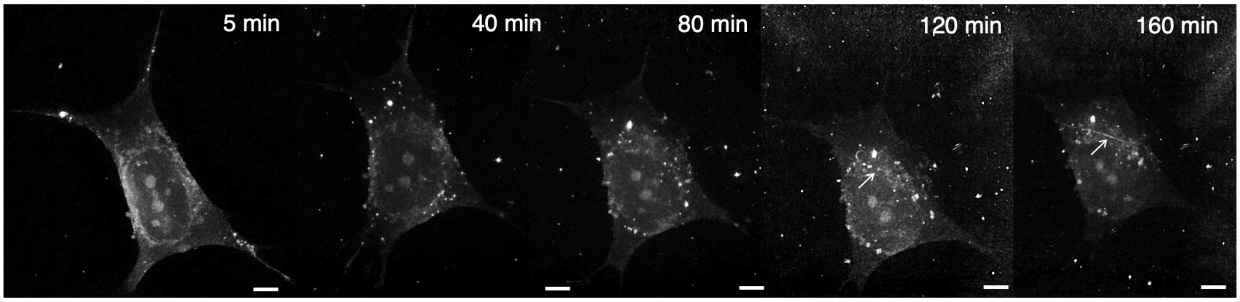


**Fig. S7. Holotomography time-lapse imaging during DTTO treatment.**

Time-course holotomography of SH-SY5Y cells exposed to 5 µg/mL DTTO for up to 160 min. DTTO monomer internalization (white dots) is detectable within the first 5 min and increases progressively during continuous exposure. At later time points, DTTO fibril begins to appear (white arrows) and elongates over time. Both monomers and fibril are clearly visible based on their refractive index (DTTO mean RI = 1.38). Scale bars: 5 µm.

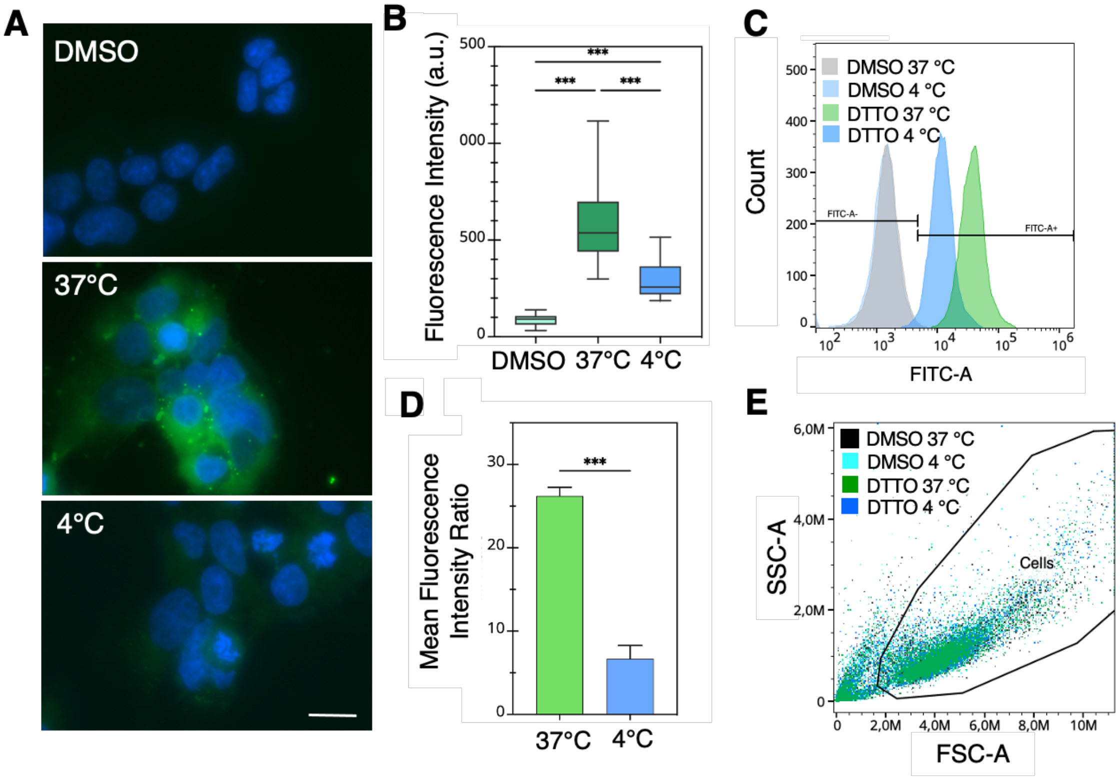


**Fig. S8. Inhibition of DTTO uptake at low temperature**

**(A)** Representative fluorescence images of SH-SY5Y cells treated with vehicle (DMSO) and DTTO 50 µg/mL at 37 ºC or 4 ºC. After treatments, cells were fixed with PFA 4%, and nuclei were stained with DAPI (blue). Scale bar: 20 µm. **(B)** Fluorescence intensities calculated from selected ROIs (n=36), each representing a cell, and corrected for the background mean; the interquartile range represent the $25^{th}$ to $75^{th}$ percentiles while the horizontal line in the middle of the box represents the median value. Whiskers represent the 5-95% percentiles. Statistical comparison was carried out using one-way ANOVA followed by Tukey's post hoc test; $***p < 0.001$. **(C)** Flow cytometry analysis: histograms of representative cell populations showing DTTO uptake at 37 ºC (green curve) and 4 ºC (blue curve). Cells were gated to identify the negative ($FITC\text{-}A^{-}$) and positive ($FITC\text{-}A^{+}$) DTTO populations. The histogram is representative of three independent experiments. Control unstained cells (DMSO) are showed as light blue and grey curves. **(D)** Mean Fluorescence Intensity ratio (MFI treated/untreated). The results represent the mean ± SD of three independent experiments. Statistical analysis was carried out using Student *t*-test; $***p < 0.001$. **(E)** Flow cytometry dot plots of vehicle-treated (DMSO) and DTTO-treated cells (50 µg/mL) at 37 ºC and 4 ºC. Cells were gated to exclude the debris (FSC-A vs. SSC-A). Plots are representative of three independent experiments.

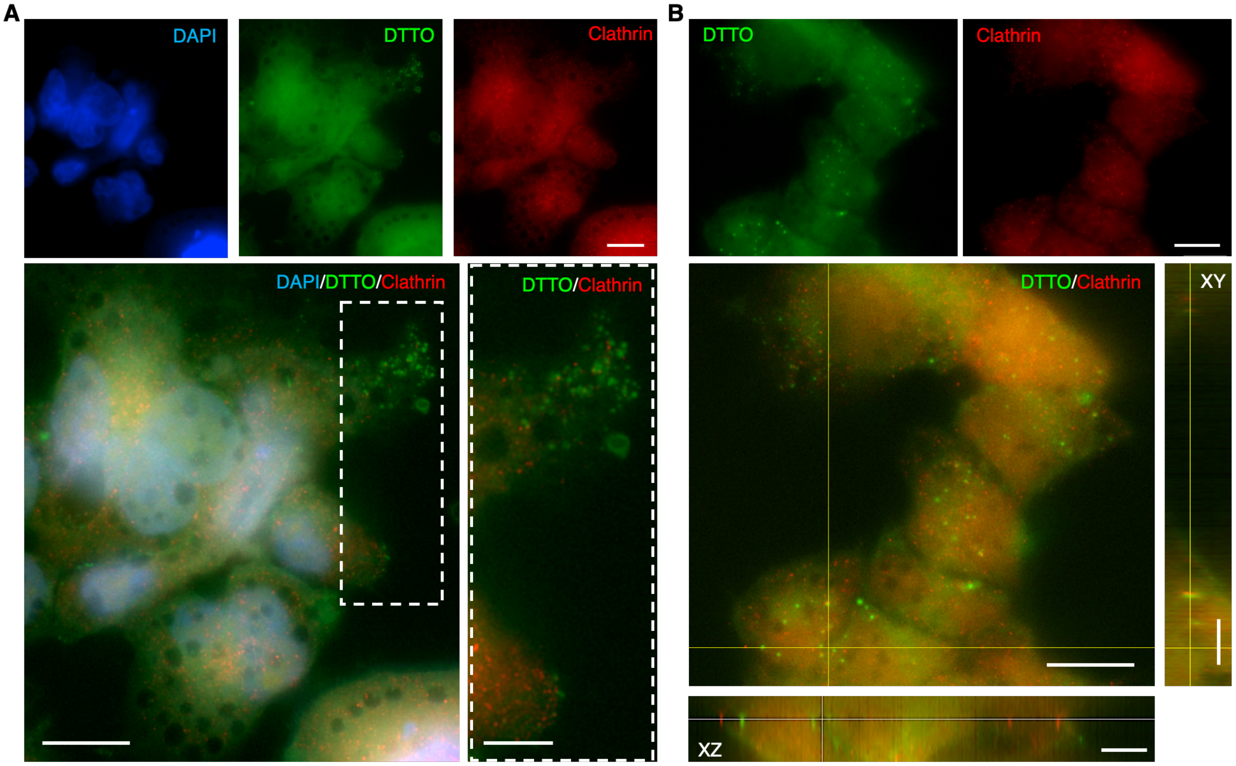


**Fig. S9. Clathrin-independent DTTO cell uptake**

**(A)** Immunofluorescence images of SH-SY5Y cells treated with 5 µg/mL of DTTO for 90 min. The magnified view, highlighted by the white dashed box, shows clear DTTO internalization. Scale bars: 20 µm, inset: 10 µm. **(B)** Single Z-plane and corresponding XZ and YZ orthogonal projections showing DTTO and clathrin signals after treating cells with DTTO 50 µg/mL for 1 hour. Scale bars: 20 µm, projection: 5 µm.

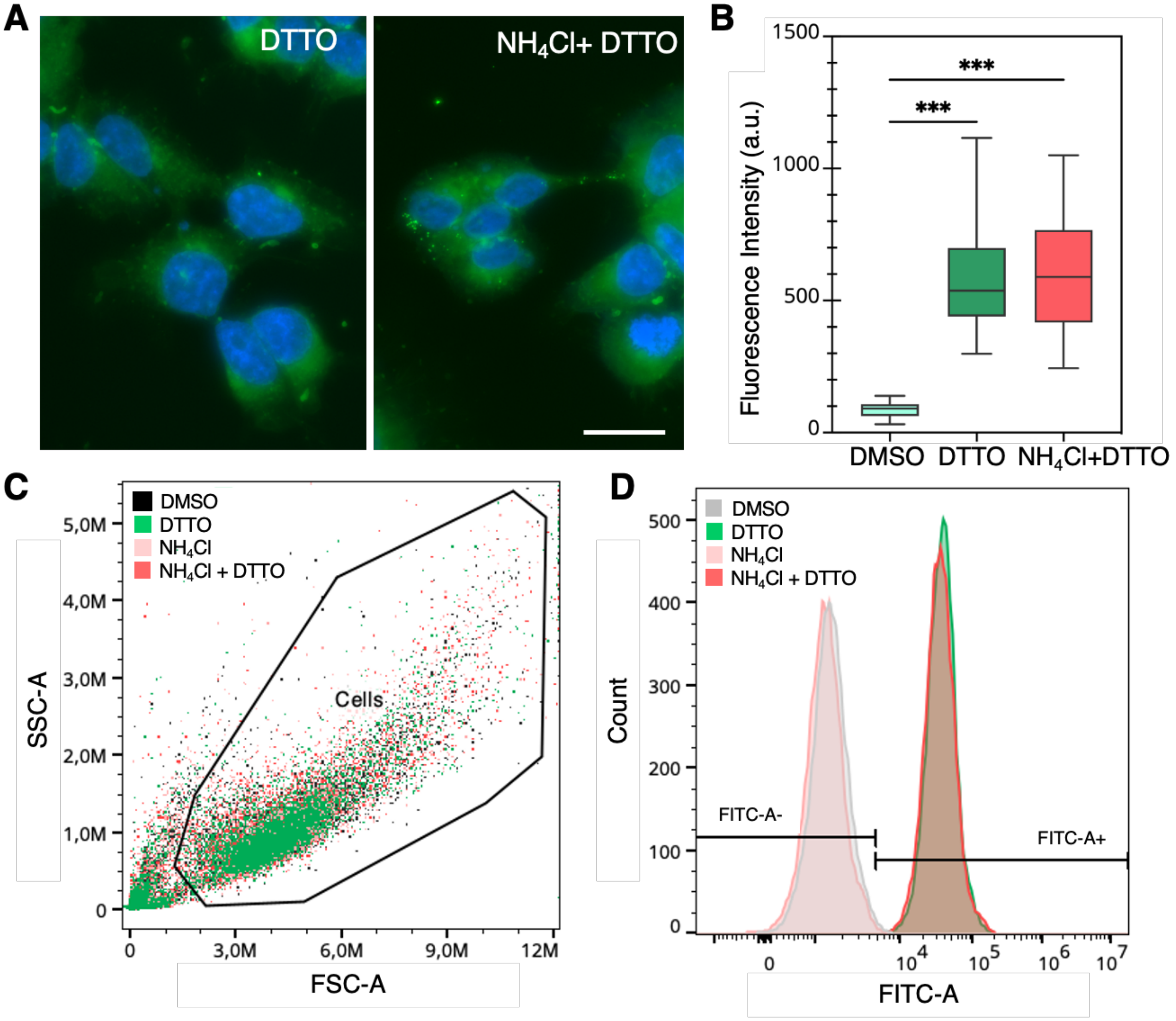


**Fig. S10. DTTO uptake is not affected by ammonium chloride**

**(A)** Fluorescence images of SH-SY5Y cells, either pre-treated with $NH_4Cl$ or not, and subsequently incubated with DTTO (50 µg/mL). After treatment, cells were fixed with PFA 4%, and nuclei were stained with DAPI (blue). Scale bar: 20 µm. **(B)** Fluorescence intensities calculated from selected cytoplasmatic ROIs (n=36), each representing a cell, and corrected for the background mean; the interquartile range represent the 25th to 75th percentiles while the horizontal line represents the median value. Whiskers represent the 5-95% percentiles. Statistical comparison was carried out using one-way ANOVA followed by Tukey's post hoc test; ***$p <$ 0.001. **(C)** Dot plots of SH-SY5Y cell populations identified by flow cytometry. Cells were gated to exclude the debris (FSC-A vs. SSC-A). Plots are representative of three independents experiments. **(D)** DTTO monomer uptake in normal condition (green curves) and with $NH_4Cl$ pre-treatment (coral curve). Cells were gated to identify the negative (FITC-$A^-$) and positive (FITC-$A^+$) populations to DTTO staining. The histogram is representative of three independent experiments. Control unstained cells (DMSO) or treated with $NH_4Cl$ are showed as grey and pink curves, respectively.

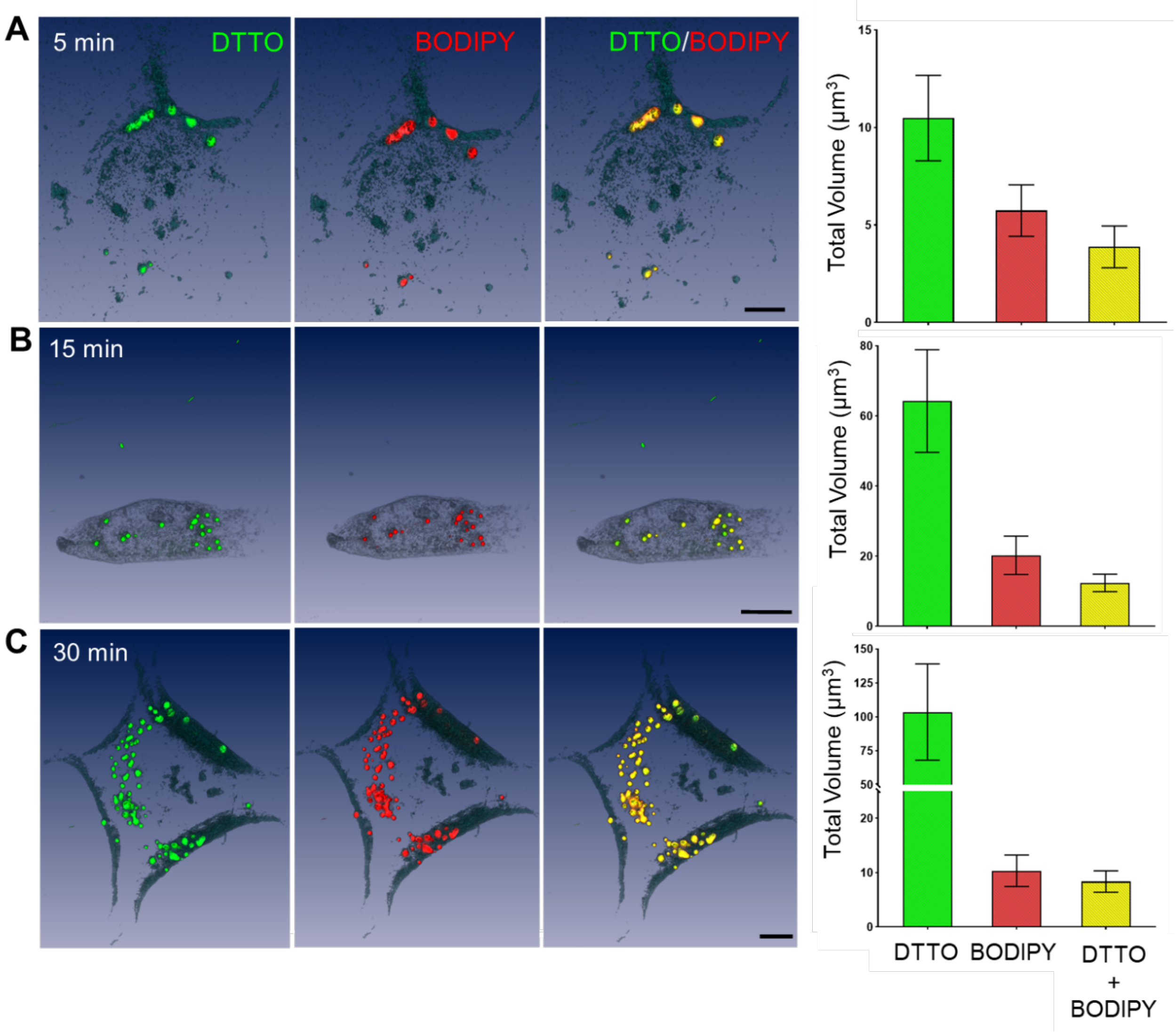


**Fig. S11. – DTTO accumulation within lipid droplets**

For each incubation time, **(A)** 5 minutes, **(B)** 15 minutes, and **(C)** 30 minutes, the left panels display the green autofluorescence signal of DTTO; the central panels show the red-channel fluorescence of BODIPY, marking lipid droplets (LDs); and the right panels illustrate in yellow the regions where the two fluorescence signals overlap. The corresponding bar graphs quantify the total fluorescence volume for DTTO, BODIPY, and their colocalization at each time point. Scale bars: 5 µm.

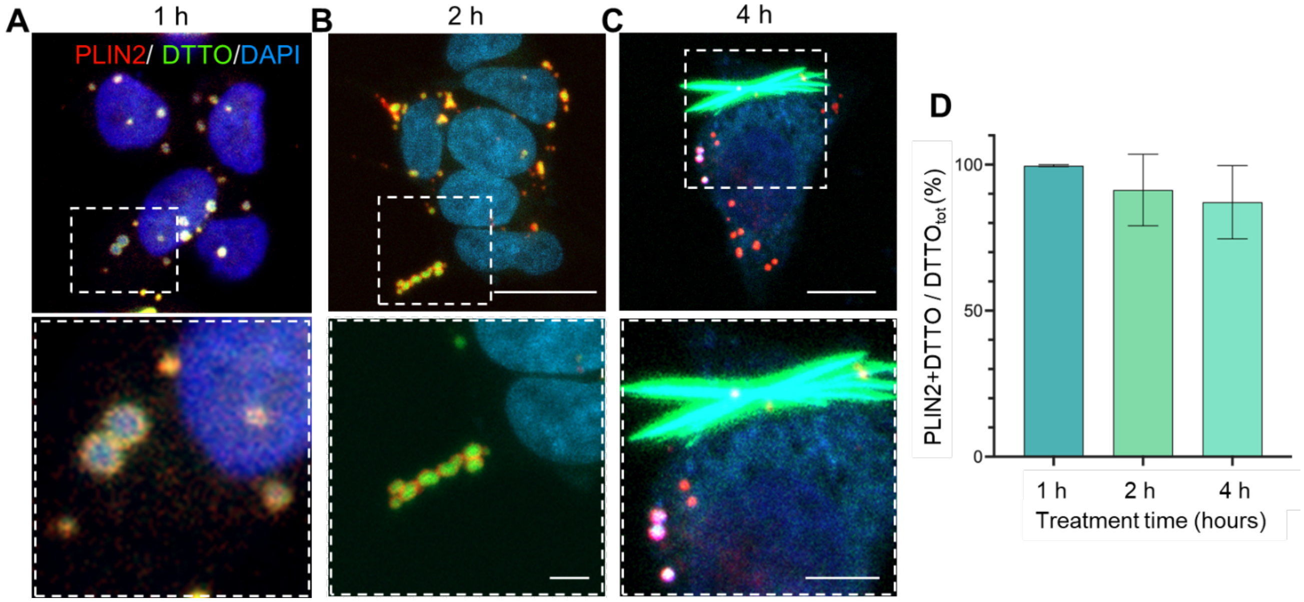


**Fig. S12. DTTO confinement in lipid droplets at increasing time**

**(A-C)** Immunofluorescence images of PLIN2-DTTO colocalization in SH-SY5Y cells treated with 5 µg/ml of DTTO for 1, 2 and 4 hours, scale bars: main 10 µm, inset 1 µm, **(C)** inset 5 µm. The dashed white boxes show magnified insets of the selected regions. **(D)** Data show the percentage of PLIN2$^{+}$, DTTO$^{+}$ over the total number of DTTO$^{+}$ vesicles (DTTO vesicles count per condition 1 h = 972; 2 h = 184; 4h = 48). Pearson's coefficient was obtained by a custom-made CellProfiler pipeline; bars represent mean ± SD.

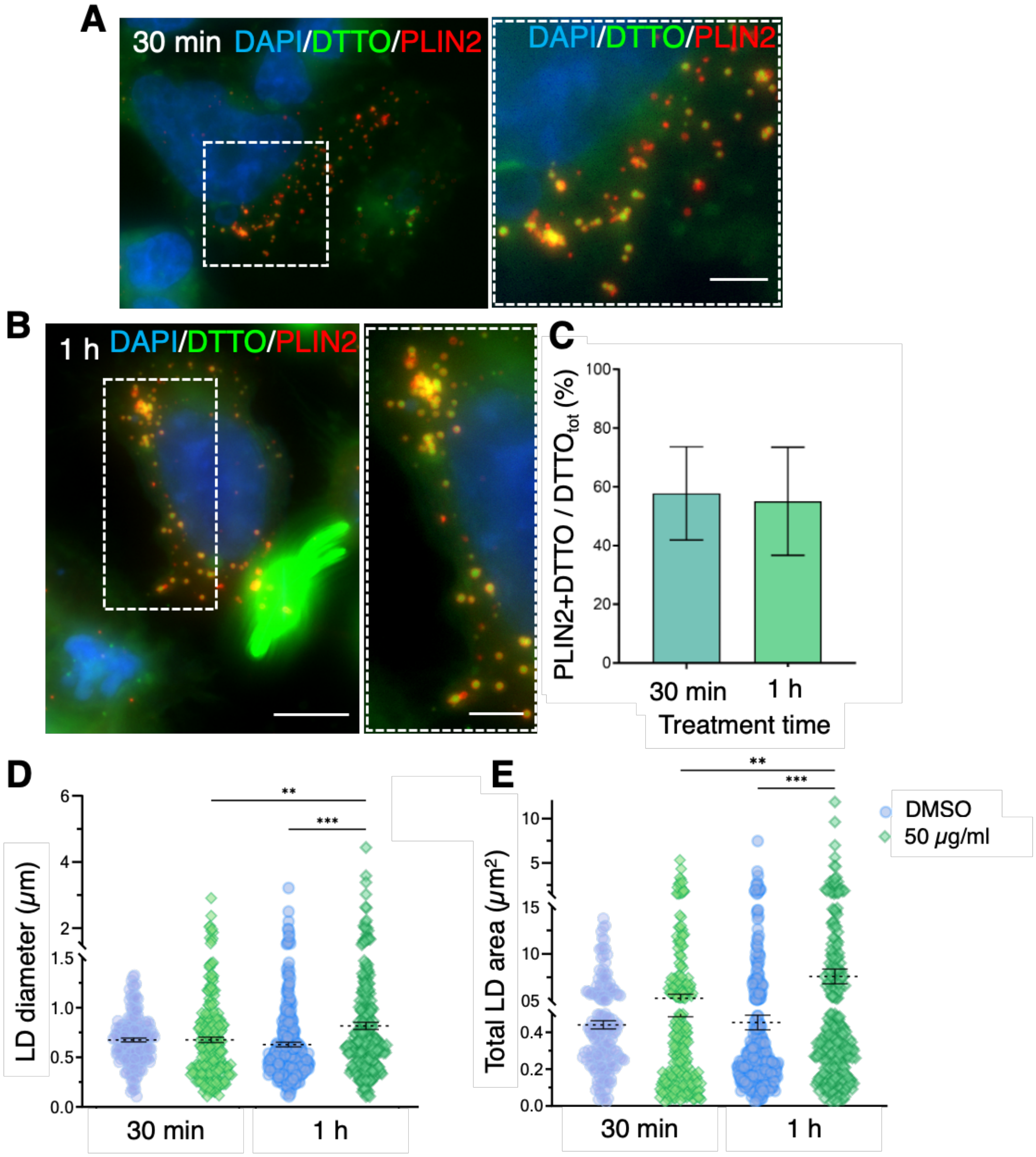


**Fig. S13. Colocalization analysis of PLIN2 and DTTO at high dose treatment**

**(A-B)** Immunofluorescence images of PLIN2-DTTO colocalization in SH-SY5Y cells treated at concentration of 50 µg/mL for 30 min and 1 hour, scale bar: 10 µm. On the right, magnified views of the areas indicated by the dashed white boxes, scale bars: 5 µm. **(C)** Data show the percentage of PLIN2$^+$, DTTO$^+$ over the total number of DTTO$^+$ vesicles (DTTO vesicles count per condition 30 min = 211; 1 h = 171). Pearson's coefficient was obtained by a custom-made CellProfiler pipeline; bars represent mean ± SD. **(D)** The diameter and **(E)** the total area of the PLIN2$^+$, DTTO$^+$ LDs were estimated and analysed by CellProfiler. Each point represents the size of a single droplet ($n_{DMSO\ 30\ min}$ = 162; $n_{DTTO\ 30\ min}$ = 248; $n_{DMSO\ 1h}$ = 287; $n_{DTTO\ 1h}$ = 256). Dashed lines represent the mean ± SEM. Statistical analysis was carried out using one-way ANOVA followed by Šídák's post hoc test; $**p < 0.01$ $***p < 0.001$.

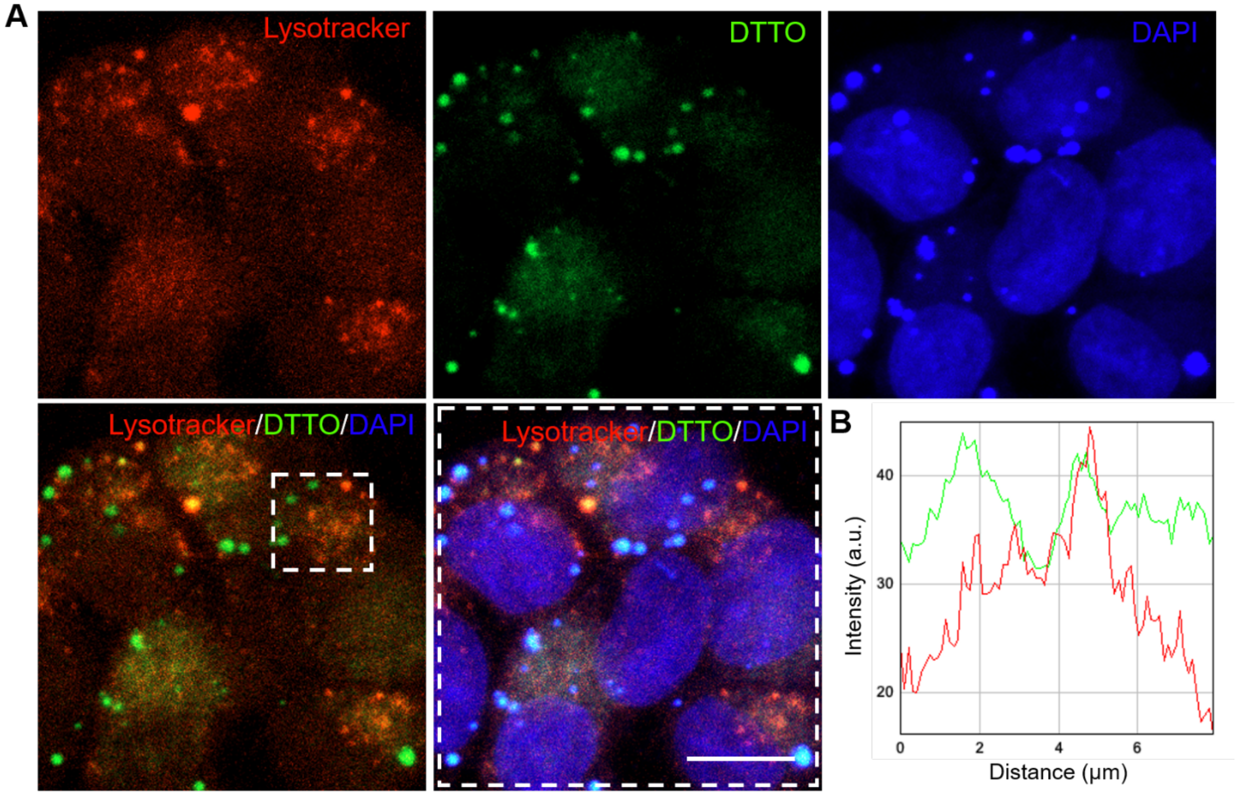


**Fig. S14. Spatial relationship between DTTO and lysosomal compartment**

**(A)** Representative fluorescence images of SH-SY5Y cells treated with 5 μg/ml DTTO and Lysotracker for 30 min. Merged images indicate that DTTO does not colocalize with Lysotracker, scale bar: 10 μm. **(B)** The line intensity profile shows the normalized fluorescence intensities of DTTO (green) and Lysotracker (red) along a line drawn through relevant regions from selected area in panel **(A)**, demonstrating only rare overlapping between the two signals.

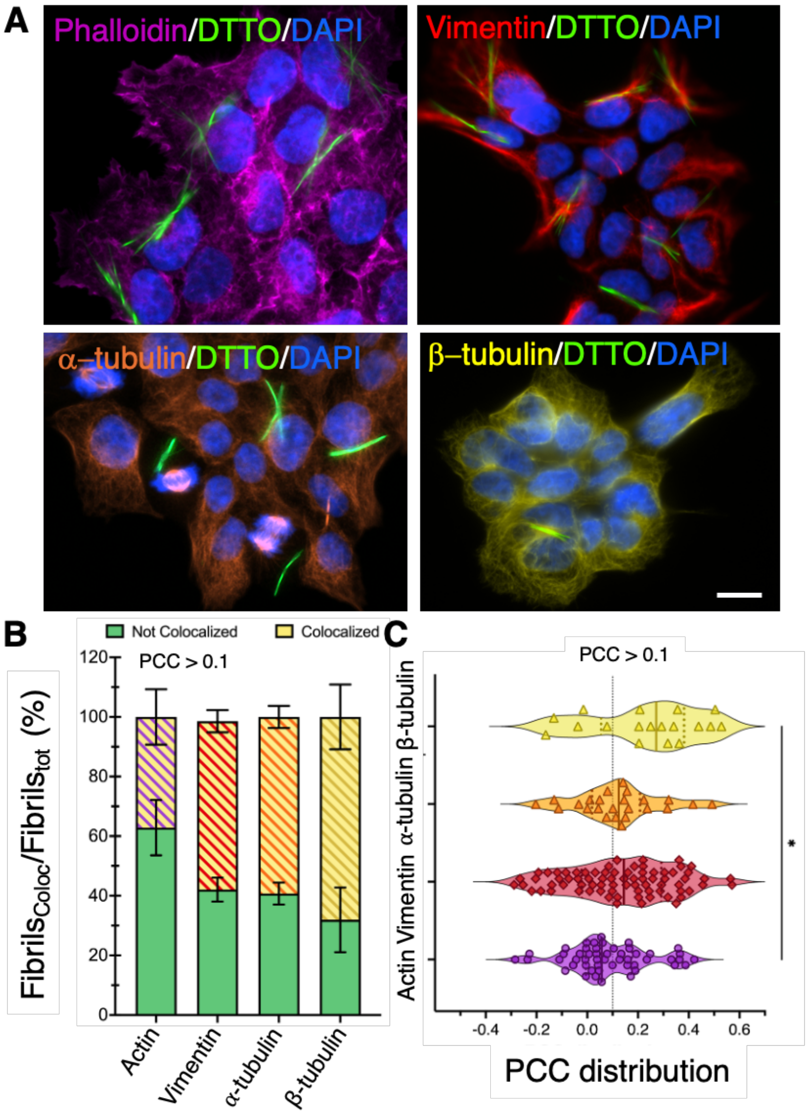


**Fig. S15. DTTO fibrils and cytoskeleton proteins correlation analysis**

**(A)** SH-SY5Y cells after 4-hour treatments with DTTO at 5 μg/mL have been fixed and incubated with anti-vimentin, anti-α and ß-tubulin antibodies and rhodamine phalloidin. Nuclei were counterstained with DAPI (blue). Scale bar: 20 μm. **(B)** Percentage of DTTO fibrils colocalizing with respective cytoskeletal proteins. Using a custom-made CellProfiler pipeline, correlation (PCC) was calculated between each fibril and the cytoskeletal protein signal; PCC values > 0.1 were considered as positive correlation (Colocalized, yellow bars), while PCC values < 0.1 were considered as negative correlation (Not Colocalized, green bars). Data represent the average percentage ± SEM obtained from a total fibril number of: phalloidin = 50, vimentin = 75, α-tubulin = 24 and ß-tubulin = 18. **(C)** Violin plot distribution of DTTO fibril PCC. Each point represents the PCC of a single fibril ($n_{actin}$= 50, $n_{vimentin}$= 75, $n_{\alpha\text{-tubulin}}$= 24, $n_{\beta\text{-tubulin}}$= 18). Solid line shows the median while dashed lines represent the upper and lower quartiles. Statistical analysis was performed using one-way ANOVA followed by Tukey's post hoc test; *$p$< 0.05.

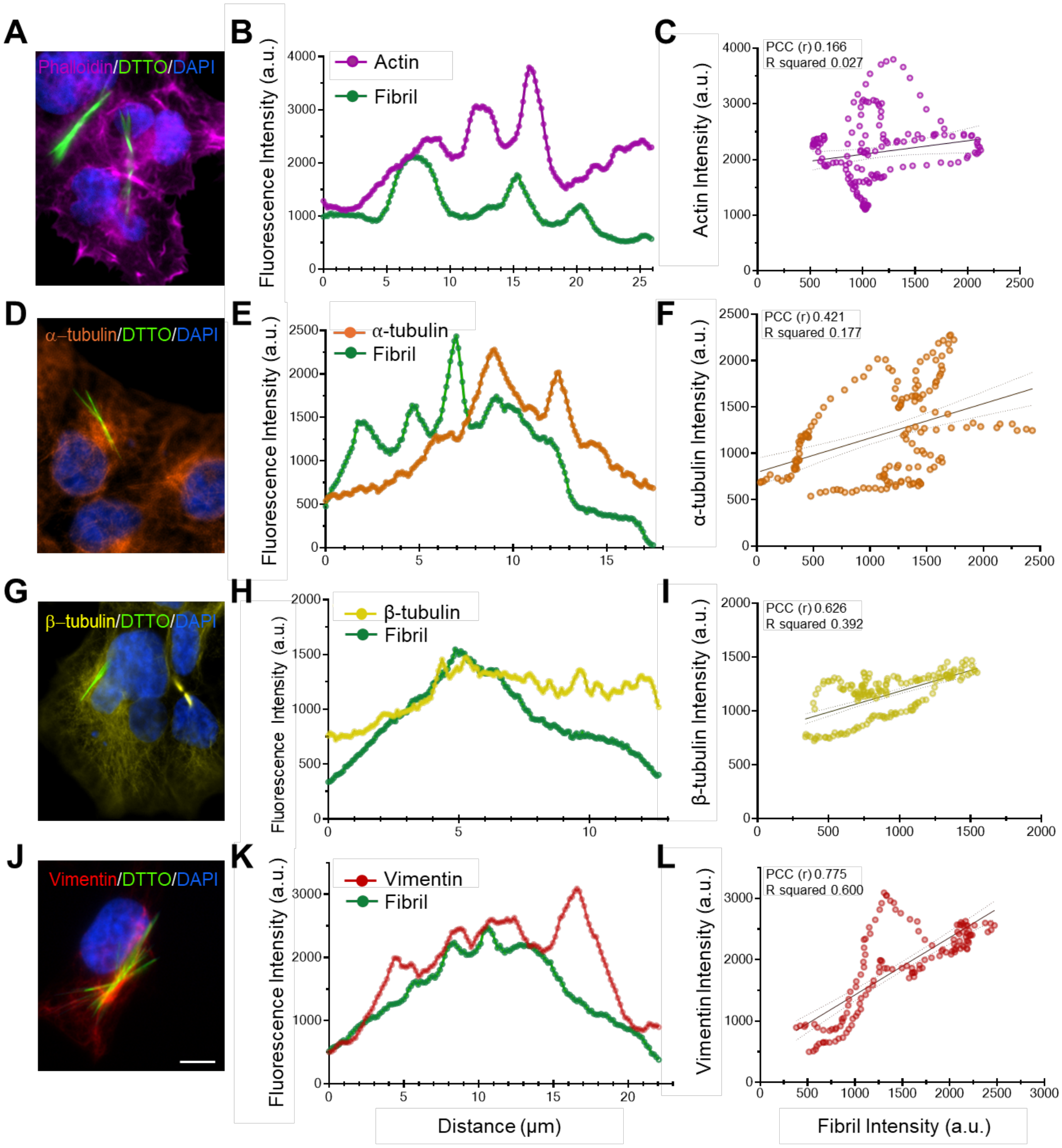


**Fig. S16. Correlation analysis on a single representative DTTO fibril**

**(A, D, G, J)** Representative immunofluorescence images of SH-SY5Y cells with DTTO fibrils synthetized after 4 h of treatment and then stained with rhodamine phalloidin, α-tubulin, β-tubulin and vimentin. Nuclei are stained with DAPI (blue). Scale bar: 10 μm. **(B, E, H, K)** Line intensity profiles of the indicated fibril obtained by Fiji software. The plot illustrates the intensity pixel distribution of DTTO (green) and actin, α-tubulin, β-tubulin and vimentin signals along the distance. **(C, F, I, L)** Scattered dot plot depicting the intensity correlation between DTTO and α-tubulin, β-tubulin and vimentin signals pixels. Each point corresponds to an individual pixel's intensity in both channels. Linear regression is overlaid as a solid line while area included between dashed lines represents 95% CI. The Pearson correlation coefficient (r) and the coefficient of determination ($R^2$) are also provided to quantify the degree of association between the two signals.

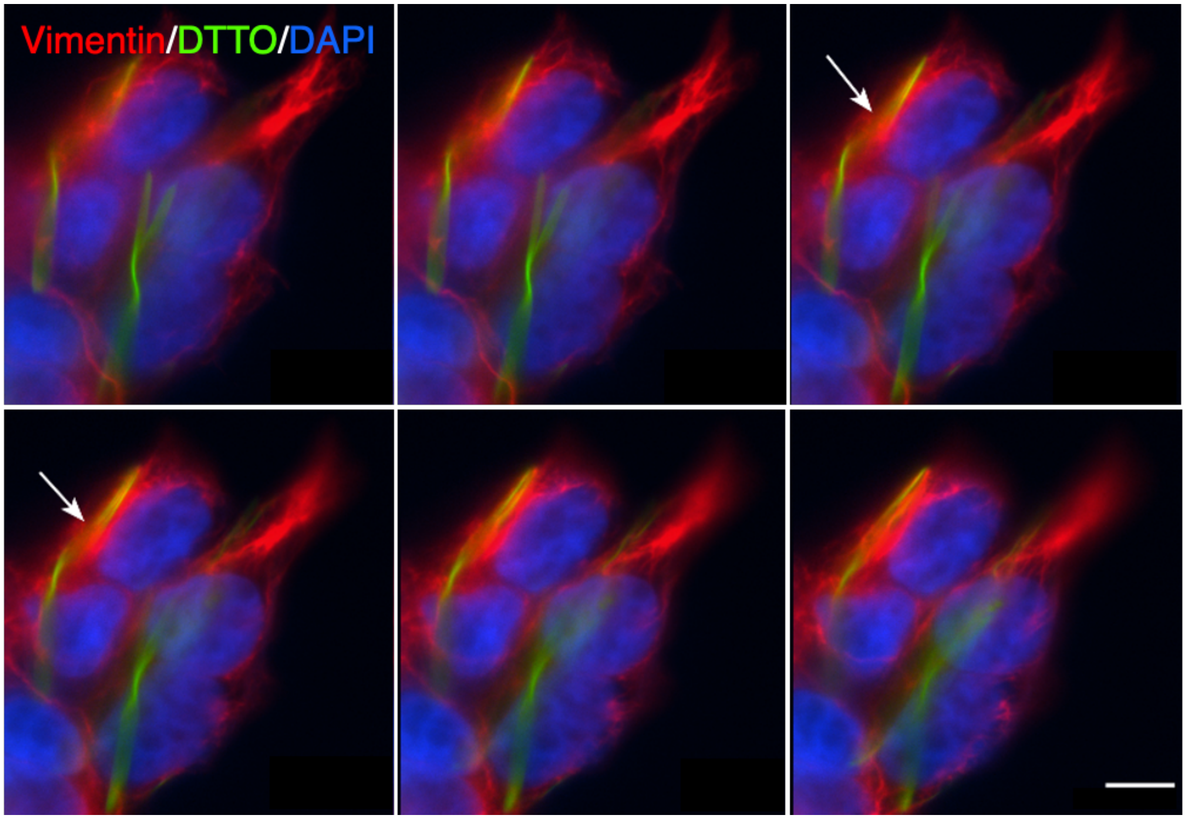


**Fig. S17. Z-stack planes montage of vimentin-stained cells producing DTTO fibrils.**

Z-stack planes at high magnification of SH-SY5Y cells stained with anti-vimentin antibody (red) after 4 hours of treatment with a low dose (5 μg/mL) of DTTO. Nuclei were counterstained with DAPI (blue), while fibrils are visible in green. White arrows indicate the DTTO fibril embedded in the complex cytoskeleton network. Scale bar: 10 μm.

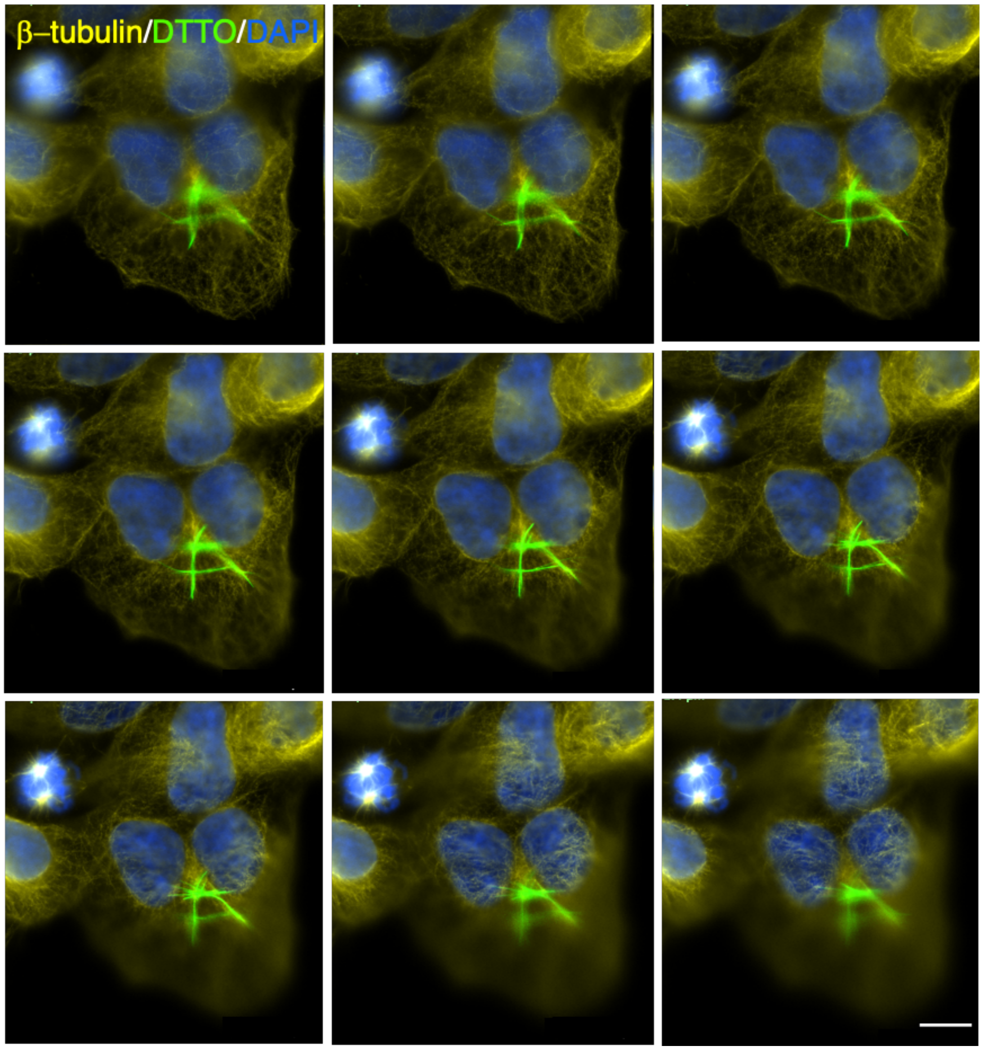


**Fig. S18. Z-stack plane montage of ß-tubulin-stained cells producing DTTO fibrils.**

Z-stack planes of SH-SY5Y cells after DTTO treatment (4 h, 5 μg/mL) and β-tubulin (yellow) immunofluorescence. Nuclei were stained with DAPI (blue), while fibrils are shown in green. Scale bar: 10 μm.

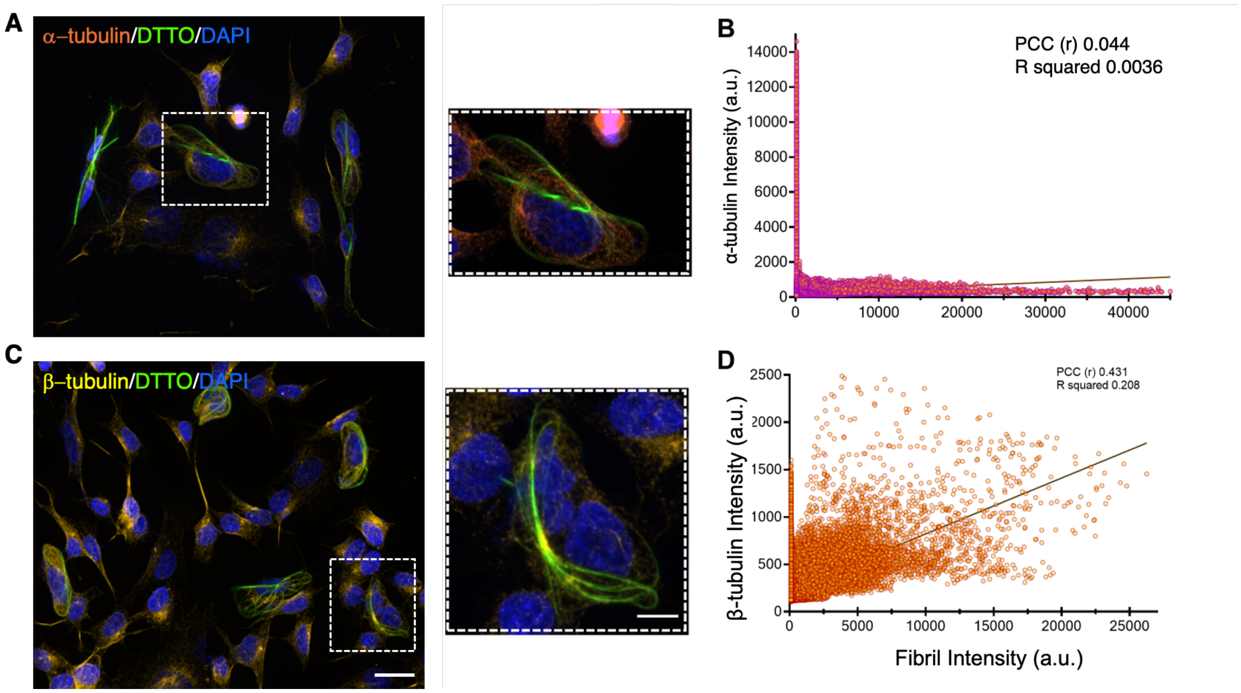


**Fig. S19. Correlation analysis of tubulins in DTTO-treated SHSY5Y cells**

SH-SY5Y cell immunofluorescence assay after overnight treatment with DTTO at 5 µg/mL. Cells were fixed, incubated with **(A)** anti-α–tubulin and **(C)** anti-β–tubulin antibodies and nuclei were labelled with DAPI (blue). Scale bars: 20 µm, insets: 10 µm. Magnified views show two representative cells with fibrils selected to assess the colocalization between DTTO, α–tubulin **(A)** and β-tubulin **(C)**. **(B-D)** Colocalization analysis was performed using JACoP plugin for Fiji software and cytofluorograms from both α-tubulin and β-tubulin were obtained.

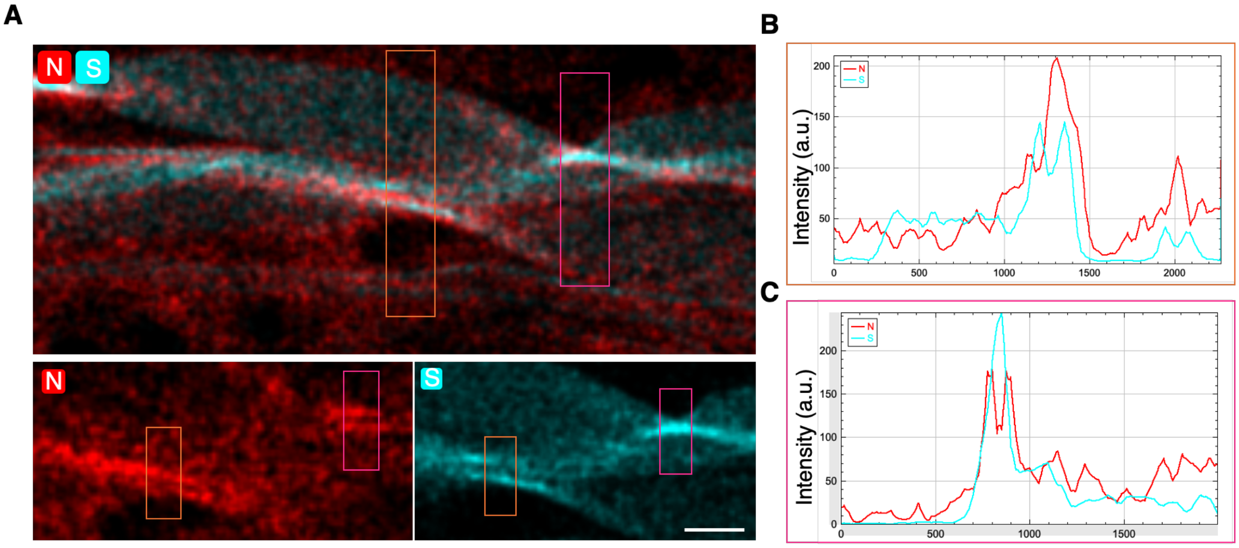


**Fig. S20. DTTO fibril isolated from SH-SY5Y cell shows helical pitches enriched in nitrogen**

**(A)** Representative elemental map obtained by EDS from an isolated fibril with Talos F200X G2 TEM/STEM operated at 200kV. Orange and magenta boxes indicate the areas selected to evaluate the intensity profile of both nitrogen and sulphur. Scale bar: 500 nm. **(B-C)** Intensity profile plots depicting the sulphur (cyan) and nitrogen (red) signals along the distance in the selected orange and magenta areas, obtained by Fiji software.

**Nano-FTIR spectroscopy**

The *IR-neaSCOPE*$^{+s}$ combines AFM with IR radiation focused on a metal-coated tip that acts as an antenna producing optical near-field interaction with the sample (*1*). The so-called IR scattering-type near-field optical microscopy (IR-s-SNOM) technique allows simultaneous access to both the morphology and the mid-IR dielectric response with resolution determined by the tip radius around 20 nm (*2*). The interferometric detection of the elastically scattered light from underneath the AFM tip provides access to amplitude and phase of the near-field scattered electric field which enables evaluating the complex dielectric response of surface and sub-surface material with spatial resolution down to the nm-scale. The real and imaginary part of the dielectric function correspond to the IR reflection and absorption properties of the material. For dielectric materials with weak Lorentz oscillators such as protein and DTTO, both amplitude and phase of the optical near-field coincide well with real and imaginary part of the complex dielectric function, and hence reflectivity and absorption in the mid-IR, giving direct access to material identification. An important aspect of the *IR-neaSCOPE*$^{+s}$ is the simultaneous demodulation at higher harmonics of the tip oscillation frequency that filters out the far-field contribution from the near-field signals and controls the penetration depth of the probed volumes (*3*).

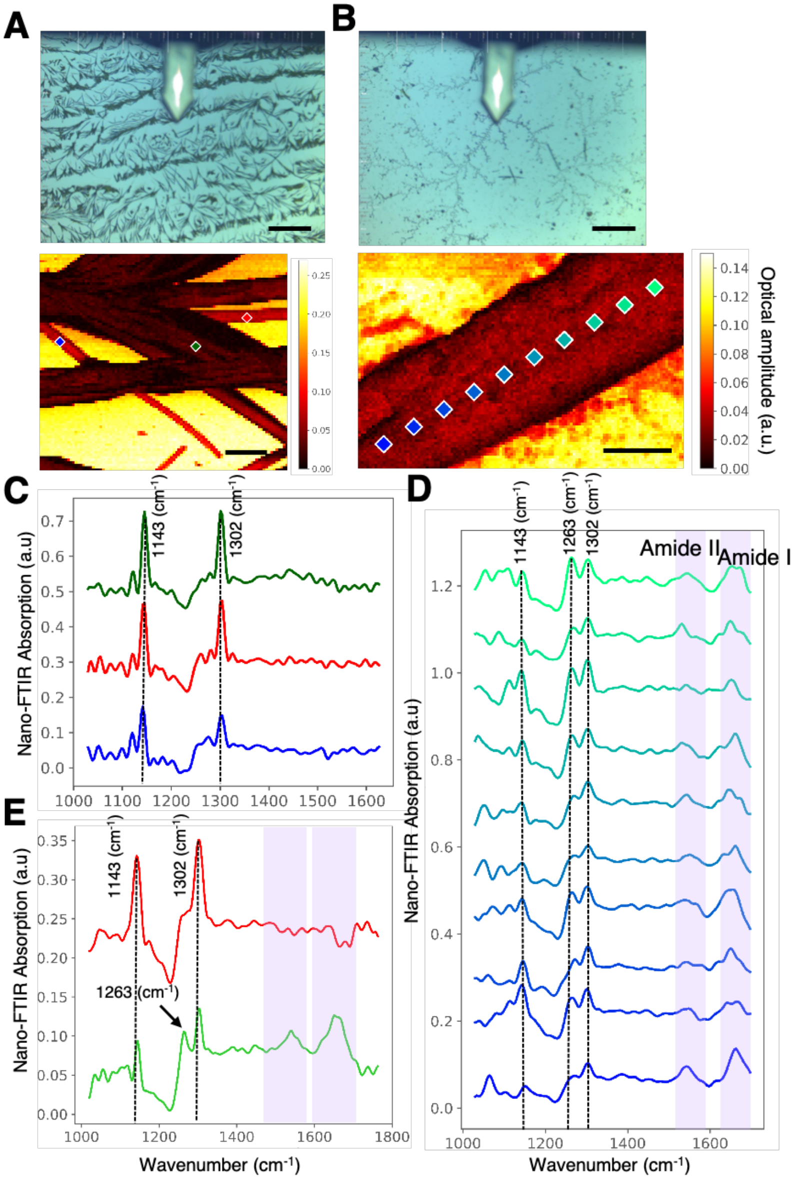


**Fig. S21. Nano-FTIR characterization of DTTO aggregates and fibrils**

Brightfield images of **(A)** DTTO aggregated in DMSO and **(B)** DTTO isolated fibril showing a restricted field of view (ca. 0.8 mm) of the whole sample, and relative reflectance images of the aggregate and fibril characterized by nano-FTIR. Coloured dots and squares indicate each single point from where spectra have been collected. Nano-FTIR spectra from **(C)** DMSO aggregate and **(D)** cell-derived fibril. Dashed lines indicate the typical bands of DTTO molecule vibrations while solid line indicates the band around 1263 $cm^{-1}$. Purple boxes indicate amide I and II characteristic bands. **(E)** Comparison between DMSO aggregate (red) and cell-derived fibril (green) nano-FTIR spectra. Each spectrum results from the average of all the detected spectra.

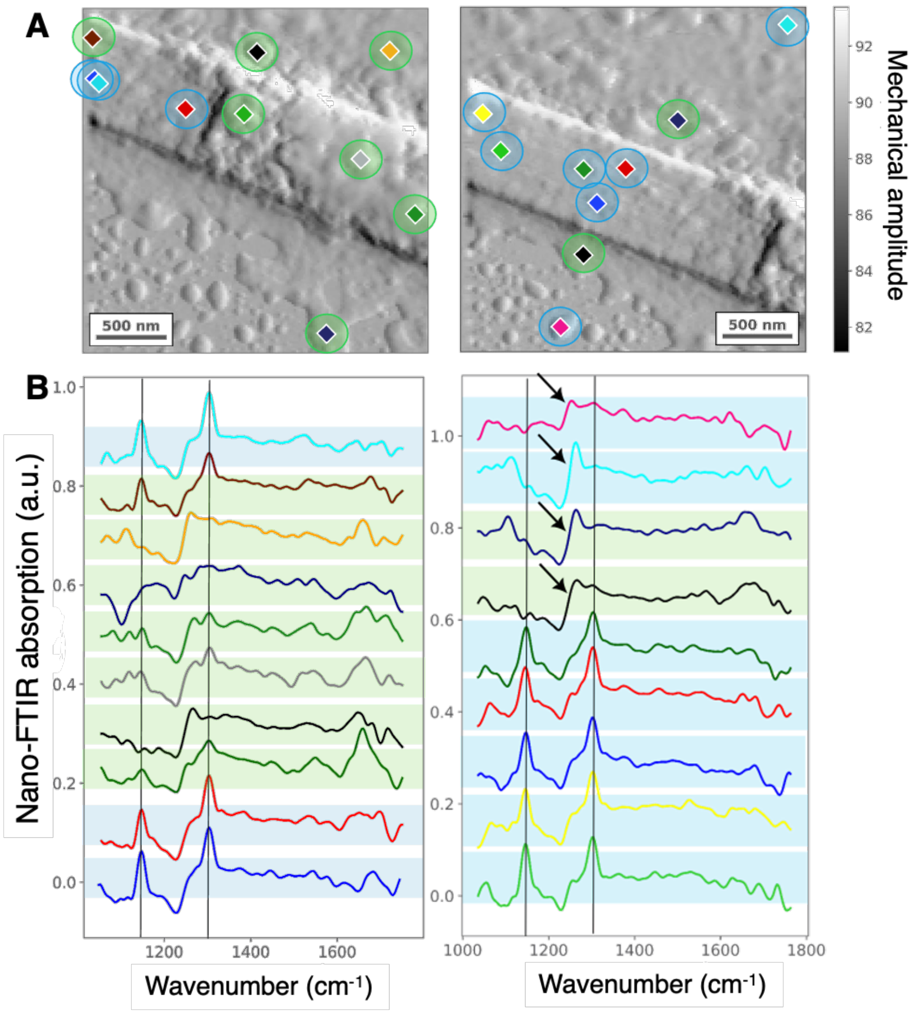


**Fig. S22. Nano-FTIR mapping of a fibril presenting heterogeneity**

**(A)** Two specific areas, one including the rupture point and the other the free-protein region, were further analysed to obtain the corresponding spectra in **(B)** from the fibril surface and the surrounding regions. Blue halos under the spectra indicate free-protein areas, while green halos indicate protein presence due to the detection of amide bands. Black arrows indicate peaks around 1263 $cm^{-1}$, suggesting the presence of a third component.

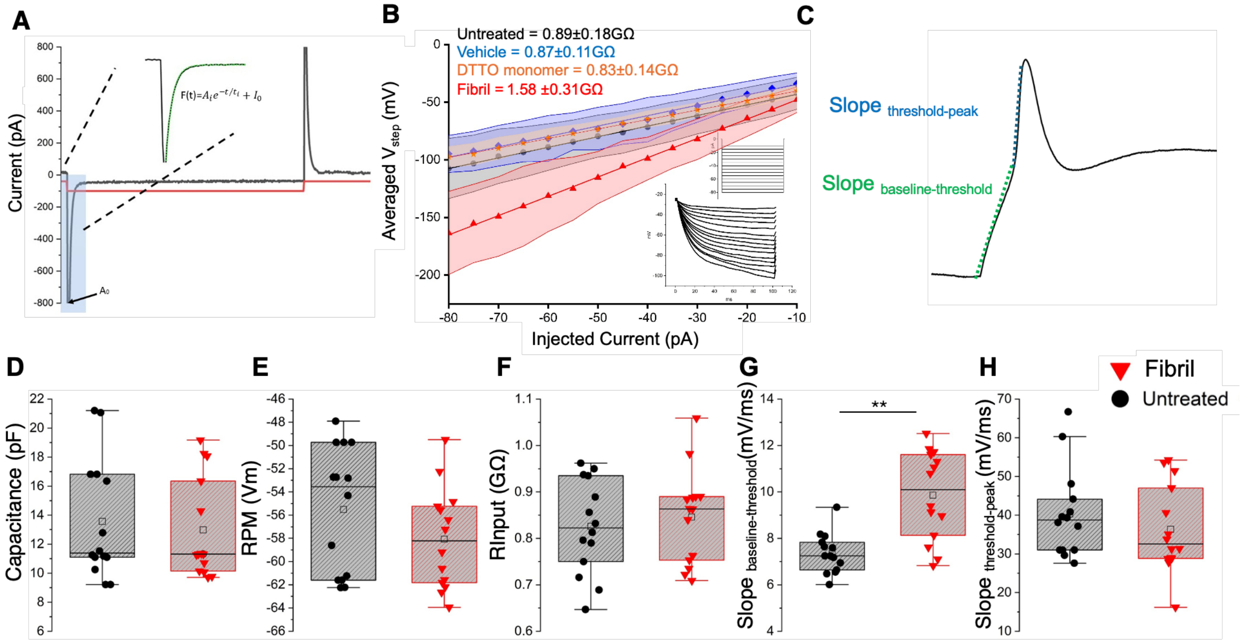


**Fig. S23. DTTO effect on RA-differentiated SH-SY5Y cells**

**(A)** Fitting approach for $C_m$ determination (see *Material and Methods*). **(B)** Comparison of the $R_{INPUT}$ estimated as the slope of a linear fit (V/I plot). **(C)** Linear regressions within different region of AP rising phase in RA-differentiated SH-SY5Y. Box plots depict mean (empty square), median (thin horizontal bar) and the 25th and 75th with whiskers showing the 5th and 95th percentile of **(D)** $C_m$, **(E)** RPM, **(F)** RInput, **(G)** Slopebaseline-threshold and **(H)** Slopethreshold-peak. One way ANOVA and Bonferroni's *post hoc* test were used for statistical comparison. $**p < 0.01$.

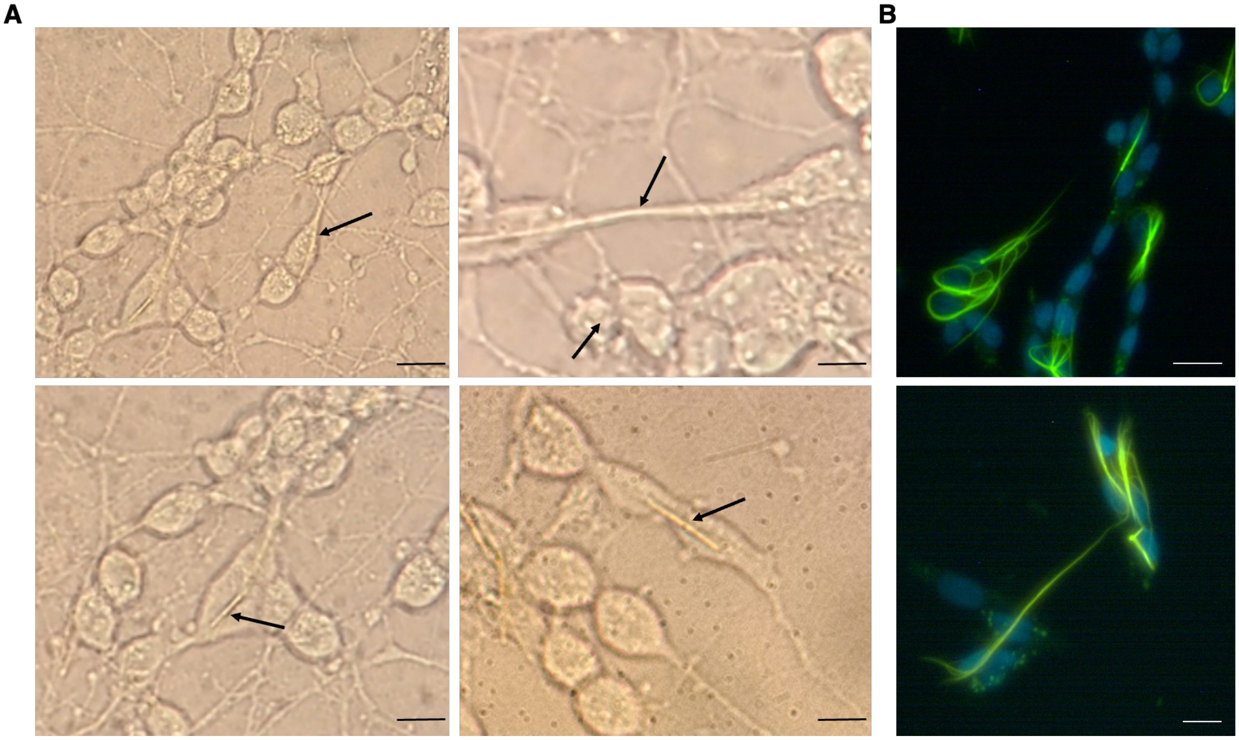


**Fig.S24. DTTO fibrils inside differentiated neuronal cells**

**(A)** Representative phase contrast and **(B)** fluorescence images of SH-SY5Y differentiated cells treated with 5 µg/mL overnight, presenting fibrils (black arrows) inside the soma and along neurites. Scale bars: **(A)** 10 µm and **(B)** 20 µm.

| Undifferentiated | $C_m$ (pF) | RPM (mV) | $R_{INPUT}$ (GΩ) |
|---|---|---|---|
| Untreated | 7.99±1.42 (n=11) | -47.72±4.6 (n=10) | 0.89±0.18 (n=13) |
| Vehicle | 8.12±1.19 (n=11) | -46.4±3.14 (n=11) | 0.87±0.11 (n=13) |
| DTTO monomer | 7.92±1.29 (n=11) | -48.54±5.53 (n=10) | 0.83±0.14 (n=11) |
| Fibril | 13.98±5.08 (n=11) | -29.27±3.46 (n=11) | 1.58±0.31 (n=14) |

**Table S1. Membrane passive properties in undifferentiated SH-SY5Y**.

Data represents the mean ± SD

| Differentiated | Untreated (n=14) | Fibrils (n=14) |
|---|---|---|
| $C_m$ (pF) | 13.56±4.11 | 12.98±3.49 |
| RPM (mV) | -55.51±5.47 | -58.07±4.32 |
| $R_{INPUT}$ (GΩ) | 0.82±0.10 | 0.84±0.10 |
| Peak (mV) | 48.93±19.69 | 33.92±18.06 |
| Time Peak (ms) | 11.74±1.82 | 8.94±2.21 |
| Slope10-90% (mV/ms) | 8.61±1.23 | 12.5±2.43 |
| Slopebaseline-threshold (mV/ms) | 7.35±0.85 | 9.86±1.93 |
| Slope1threshold-peak (mV/ms) | 40.32±11.52 | 36.35±11.38 |

**Table S2. Membrane passive properties and AP electrophysiological parameters in differentiated SH-SY5Y**

Data represents the mean ± SD.

**Movies S1–S2.** 3D reconstruction of DTTO fibrils inside SH-SY5Y cells treated for 24 hours with 5 µg/mL DTTO, imaged by super-resolution microscopy. β-tubulin is shown in yellow, nuclei in blue and DTTO fibrils in green.

**Movies S3–S5.** Merged fluorescence and holotomography 3D reconstructions along the y-axis of SH-SY5Y cells treated with CellMask (red, labeling plasma membrane lipids) and 5 µg/mL DTTO (green) for 40 minutes, 4 hours, and 24 hours before fixation. Nuclei are shown in grey (RI = 1.362–1.367).

**Movies S6–S8.** Merged fluorescence and holotomography Z-planes of SH-SY5Y cells treated with CellMask (red, labeling plasma membrane lipids) and DTTO (green) for 40 min, 4 hours, and 24 hours before fixation. Nuclei are shown in grey (RI = 1.362–1.367).

### References


1. B. Knoll, F. Keilmann, Near-field probing of vibrational absorption for chemical microscopy. *Nature* **399**, 134-137 (1999).
2. F. Huth *et al.*, Nano-FTIR absorption spectroscopy of molecular fingerprints at 20 nm spatial resolution. *Nano Lett* **12**, 3973-3978 (2012).
3. L. Mester, A. A. Govyadinov, S. Chen, M. Goikoetxea, R. Hillenbrand, Subsurface chemical nanoidentification by nano-FTIR spectroscopy. *Nat Commun* **11**, 3359 (2020).
